# Biochemical and structural insights into the auto-inhibited state of Mical1 and its activation by Rab8

**DOI:** 10.1101/2024.06.17.599268

**Authors:** Amrita Rai, Petra Janning, Ingrid R. Vetter, Roger S. Goody

## Abstract

Mical1 regulates F-actin dynamics through the reversible oxidation of actin, a process controlled by its interactions with various proteins. Upon binding to Rab8 family members, Mical1 links endosomes to the cytoskeleton, promoting F-actin disassembly. In the absence of Rab, Mical1 exists in an auto-inhibited state, but its biochemical characterization remains incomplete. Our study reveals that the N-terminal MO-CH-LIM domains of Mical1 form an intramolecular complex with its C-terminal bMERB domain. Mutational analysis, guided by the AlphaFold2 model, identifies critical residues at the binding interface. Additionally, we demonstrate that full-length Mical1 binds to Rab8 in a 1:2 stoichiometry, thereby releasing auto-inhibition. Through structure-based mutational studies, we uncover allostery between the N and C-terminal Rab binding sites. Notably, Rab binding at the high-affinity C-terminal site precedes binding at the N-terminal site, suggesting a sequential binding mode. These findings elucidate how Rab8 binding releases the MO-CH-LIM domains from the Mical1 bMERB domain, facilitating interactions with other proteins and the actin cytoskeleton, thereby modulating actin dynamics.

## Introduction

Actin dynamics is a highly controlled process, and the precise assembly and disassembly of actin is necessary for many cellular processes, including motility, adhesion, morphogenesis, and cytokinesis^1–3^. Actin-binding proteins (ABPs) are known to control filament polymerization (F-actin), its organization (bundling), and actin disassembly (G-actin). Apart from ABPs, actin post-translational modifications such as phosphorylation, oxidation, acetylation, arginylation, SUMOylation, ubiquitination, and others, regulate actin dynamics, fine-tune actin organization, and its interaction with ABPs^4^. The Molecule Interacting with CasL (Mical) protein family members include Mical1, Mical2, and Mical3, which are monooxygenases that oxidize actin and regulate cytoskeleton organization in the presence of the coenzyme NADPH and of O ^5^. Mical1 specifically oxidizes Met44 and Met47 of the DNaseI-binding loop (D-loop) in actin monomers^6^. These residues are involved in bridging actin subunits, thereby regulating F-actin stability^6^. Methionine oxidation leads to F-actin depolymerization, a process that can be selectively reversed by methionine sulfoxide reductases. For example, MsrB reduces Met44 and Met47 of oxidized monomeric actin (G-actin), leading to the repolymerization of actin^7,8^. Mical proteins play a crucial role in actin assembly regulation during muscle organization, axon guidance, drosophila bristle development, cell shape, cell viability, cardiovascular integrity, vesicular trafficking, and regulation of nuclear actin^9^. Mical1 has been implicated in several different cancers, including pancreatic^10^, breast^11^, gastric^12^, colorectal^13^, and melanoma^14^ as well as in neurological and mental health disorders such as neurodegeneration^15^, spinal cord injury^16^, and epilepsy^17^.

Mical family members contain an N-terminal monooxygenase (MO) domain which plays a crucial role in actin-depolymerization, a calponin homology (CH) domain, a LIM (Lin-11, Isl-1, and Mec-3) domain which provides a protein-protein interaction interface, and a C-terminal bMERB domain that interacts with Rab proteins^5,18^. Apart from these domains, Mical family members have a proline-rich region that interacts with SH3 domain-containing proteins^19^. Mical1 plays a crucial role in synapse development^20^. Mical1 and Mical3 are required for cytokinesis and vesicular trafficking. They regulate these processes by regulating F-actin dynamics by disassembling the actin filaments. Mical-like (MICAL-L) family members lack the MO domain and play critical roles in vesicular trafficking^21^. Humans have two Mical-like proteins (Mical-L1 and Mical-L2/JRAB) whereas Drosophila has a single Mical and a single Mical-like protein. Several studies have reported that in the absence of stimuli, the Micals and Mical-like family members exist in an auto-inhibited state^9^. However, biochemical, and structural characterization of the auto-inhibited state is missing.

Rab GTPases are players that have a crucial role in vesicular trafficking and more than 60 Rabs have been reported for humans^22^. Similar to other small GTPases, they act as molecular switches, switching between GTP-bound active state and GDP-bound inactive state^22^. The switching process is spatiotemporally regulated by guanine nucleotide exchange factors (GEFs) which facilitate the nucleotide exchange from GDP to GTP and their GTPase activity is enhanced by GTPase-activating proteins (GAPs)^23–25^. In the active GTP bound state, Rabs interacts with specific effector molecules to execute distinct downstream functions^22^. MICAL family members are one such Rab8 family effector. It has been shown that the binding of Rab molecules activates these family members by releasing auto-inhibition^26,27^. We have previously shown that in the case of some family members, such as Mical1/3 and EHBP1L1, Rab binds to the bMERB domain in a 1:2 stoichiometry, whereas other bMERB family members have only a C-terminal high affinity binding site^28–30^. However, it is still to be shown whether full-length Mical1 binds to two Rab molecules and if there is any allosteric interaction between the two binding sites.

In this study, we comprehensively characterized the auto-inhibited and active states of human Mical1 using biochemical and structural methods. Via isothermal titration calorimetry (ITC), we confirmed that the N-terminal MO-CH-LIM domains interact with the C-terminal bMERB domain, requiring the full-length bMERB domain for this interaction. Crosslinking-mass spectrometry (XL-MS) experiments supported the auto-inhibited state of Mical1. Utilizing AlphaFold2^31^, we constructed a model revealing the auto-inhibited state and identified crucial interacting residues. Additionally, we confirmed the 1:2 stoichiometry of Rab8a binding to full-length Mical1. Next, we investigated whether allostery exists in the N- and C-terminal Rab binding sites. To answer this, we have used the bMERB domain, since it shows similar Rab affinity as full-length Mical1. Biochemical experiments show that Rab8 first binds to the high-affinity C-terminal site, facilitating subsequent second Rab8 binding to the low-affinity N-terminal site. These findings underscore the essential role of Rab binding in Mical1 activation.

## Results and Discussion

### Mical1 exists in an auto-inhibited state

Human Mical1 comprises four domains: an N-terminal monooxygenase (MO), a central calponin (CH), a LIM (Lin-11, Isl-1, and Mec-3) domain, and a C-terminal bivalent Mical/EHBP Rab-binding (bMERB) domain. Domain boundaries are taken from isoform 1 (UniProt ID: Q8TDZ2) (Supplementary Fig. 1). To characterize the auto-inhibited and active states, we purified full-length Mical1 and various deletion constructs (Fig. 1a and Supplementary Table. 2). All constructs except the bMERB domain are tagged with a C-terminal histidine tag. Further, to determine the oligomeric states of the proteins, their native masses were determined by size exclusion chromatography combined with multi-angle light scattering analysis (SEC-MALS). Results showed that all the purified constructs predominantly exist as monomers. The average molar masses were as follows: full-length Mical1, 112.2 kDa (theoretical mass: 118.94 kDa); MO-CH-LIM (residues 1-781), 86.2 kDa (theoretical mass: 86.48 kDa); MO-CH (residues 1-615), 69.4 kDa (theoretical mass: 68.23 kDa); MO (residues 1-488), 55.78 kDa (theoretical mass: 55.14 kDa); bMERB (residues 918-1067), 18.13 kDa (theoretical mass: 17.83 kDa) (Fig. 1b-f).

**Fig. 1:**
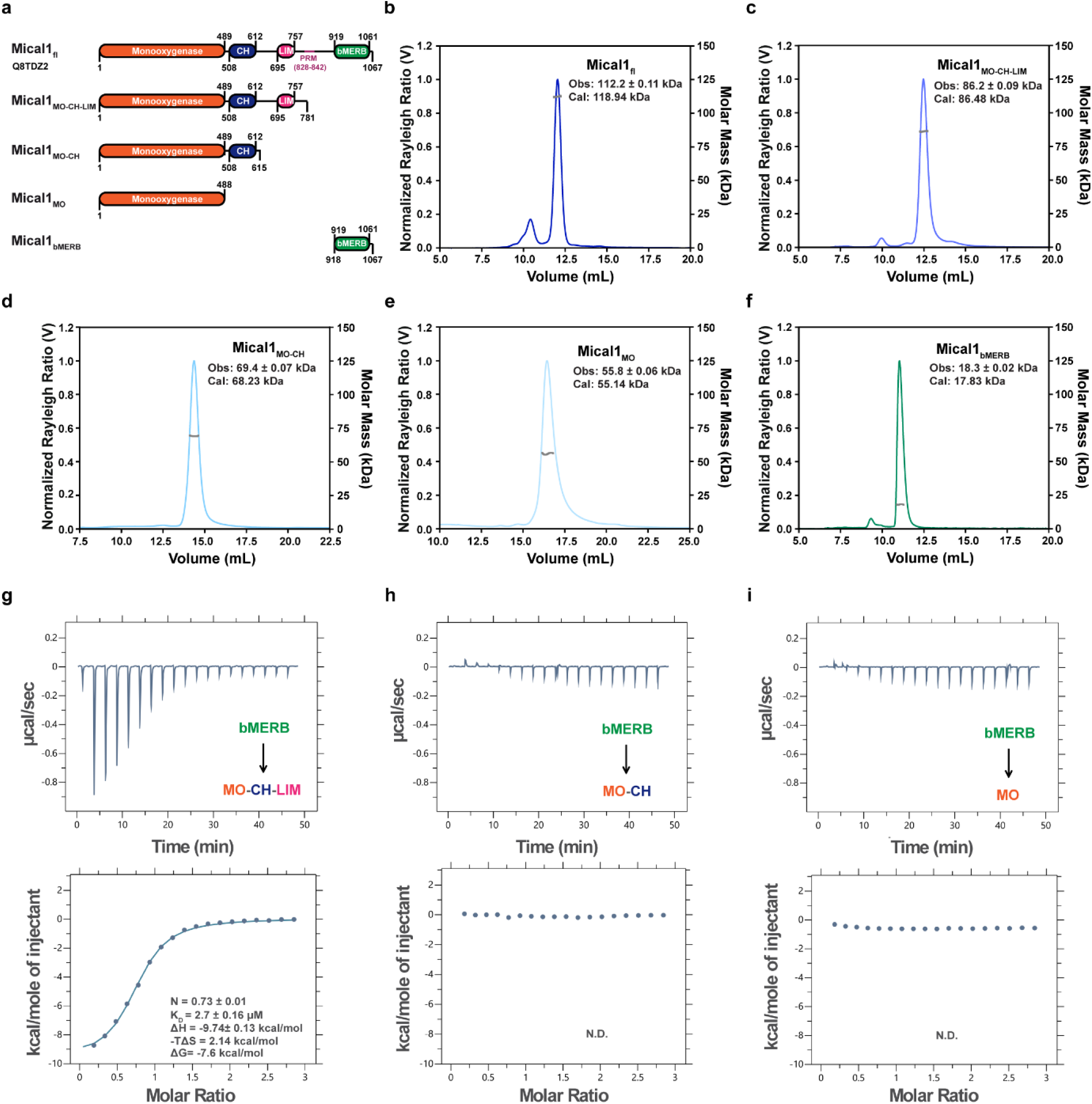
Mical1 MO-CH-LIM domains form an intramolecular complex with its bMERB domain. **(a)** A schematic diagram of human Mical1: Mical1 contains an N-terminal monooxygenase (MO) domain, calponin homology (CH) domain, Lin11, Isl-1, and Mec-3 (LIM) domain, and a C-terminal coiled-coil bMERB (bivalent Mical/EHBP Rab binding) domain. PRM: proline-rich motif. (**b-f**) Size exclusion chromatography (Superdex 200) coupled with multi-angle light scattering (SEC-MALS) was performed for full-length Mical1, MO-CH-LIM, MO-CH, and MO, while size exclusion chromatography (Superdex 75) coupled with SEC-MALS was conducted for the bMERB domain. The Rayleigh scattering is represented in blue, and the molecular weight distribution across the peak is shown in gray. The measured molecular weight closely corresponds to that of a monomer. (**g-i)** MO-CH-LIM domain construct interacts with the bMERB domain. Binding affinities were measured by titrating the bMERB domain (600 µM) with the MO-CH-LIM/MO-CH/MO domain (40 µM). Integrated heat peaks, corrected for the offset by subtracting the control experiment, were fitted to a one-site-binding model, yielding the binding stoichiometry (N), enthalpy (ΔH), entropy (ΔS), and dissociation constant (K_D_). Clear complex formation was observed for the MO-CH-LIM, whereas no complex formation was observed for the MO-CH and MO. The data represents results from at least three independent experiments. N.D. denotes not detected.

Several studies have indicated that, in the absence of Rab, the bMERB domain forms an intramolecular interaction with the other domains (CH/LIM/MO-CH-LIM)^26,32–35^. Full-length Mical is catalytically inactive, with its enzyme activity somehow inhibited by the C-terminal domain^36^. Fremont *et al*., proposed a Mical1 auto-inhibition model based on F-actin depolymerization assays and cell biology data, suggesting that the N-terminal monooxygenase and CH and LIM domains (MO-CH-LIM) interact with the bMERB domain to form an inactive complex^26^. The binding of Rab releases auto-inhibition, resulting in F-actin depolymerization^26,36,37^. However, the biochemistry of the auto-inhibited state is not well understood. Therefore, we aimed to confirm and quantify the intramolecular interaction between the N- and C-terminal domains of Mical1 by investigating the interaction between the separately purified C-terminal bMERB domain and the N-terminal MO/MO-CH/MO-CH-LIM domains via ITC. We observed binding between the MO-CH-LIM domain construct and the bMERB domain, driven by enthalpy with a K_D_ value of 2.7 ± 0.16 µM and a 1:1 stoichiometry. However, no binding was observed between the isolated bMERB domain and the MO-CH/MO domains (Fig. 1g-i). Our previous work also showed that the Mical1 CH-LIM construct and isolated CH and LIM domains do not interact with the bMERB domain^30^. Thus, our data indicate that all three domains (MO-CH-LIM) are necessary for stable bMERB interaction (Fig. 1g-i). Like Mical1, other F-actin interacting proteins, such as Myosin V, Talin2, and Vinculin, also exist in an auto-inhibited state, with their activation being spatially and temporally regulated^38,39^.

### Full-length bMERB domain is essential for MO-CH-LIM interaction

Mical1 and Mical3, among other bMERB family members, feature a high-affinity C-terminal Rab-binding site (Ct-RBS2; helices 2-3) and a lower-affinity N-terminal site (Nt-RBS1; helices 1-2). In contrast, EHBP1L1’s bMERB domain has two Rab-binding sites of similar affinities^28,29^. EHBP1, another family member, has a single Rab-binding site and forms an auto-inhibited complex where helices 1-2 of the bMERB domain constitute the CH binding site, with Rab binding at the C-terminal site (helices 2-3) releasing auto-inhibition^30^. To pinpoint the exact binding site on Mical1, we generated two deletion constructs of its bMERB domain, deleting either the N-terminal or C-terminal helix, and tested their interaction with MO-CH-LIM. Our ITC data confirm that the full-length bMERB domain is necessary for a stable interaction, as the deletion constructs did not form a stable complex. (Fig. 2a-c).

**Fig. 2:**
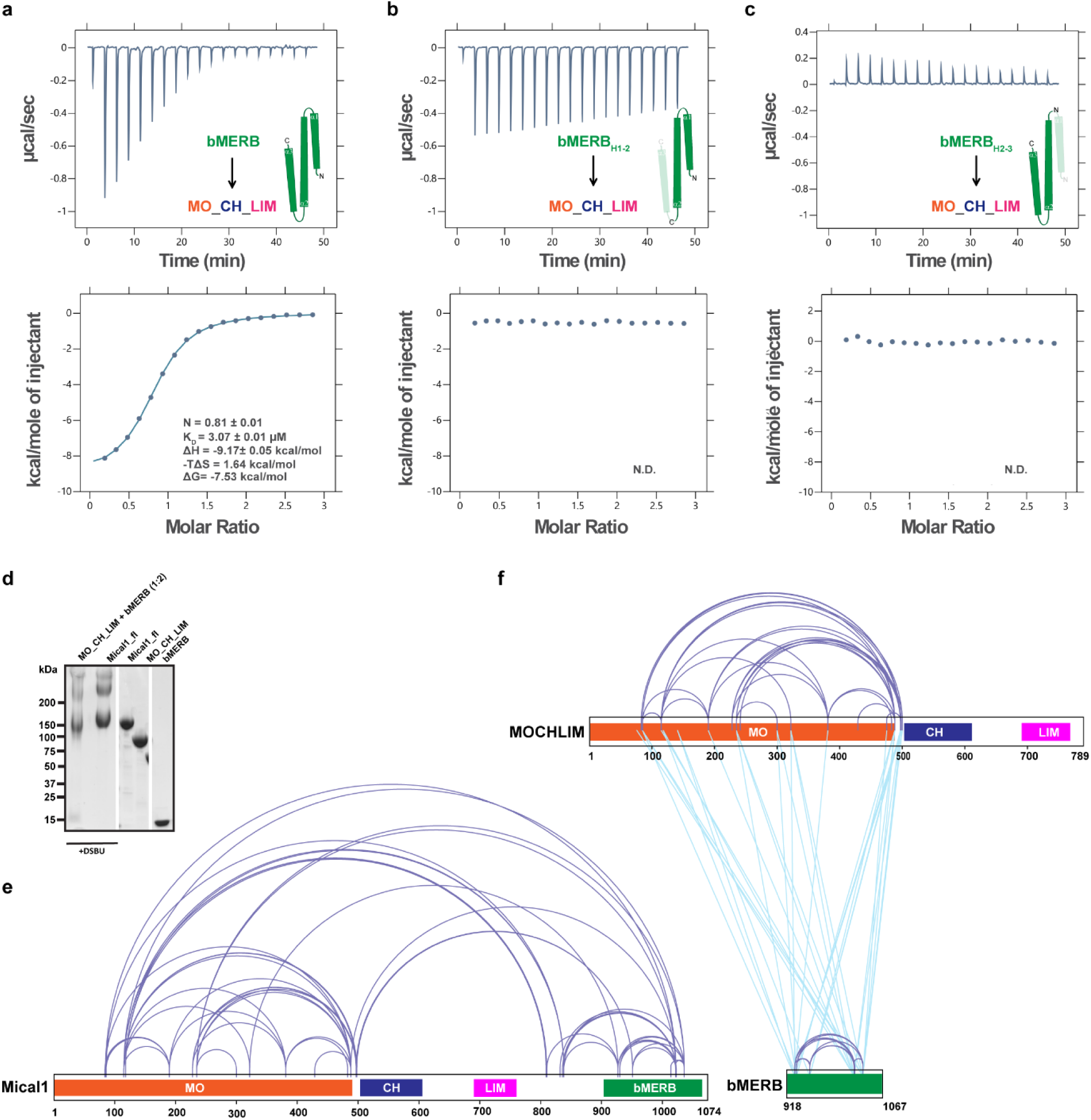
Full-length bMERB domain is essential for interaction with MO-CH-LIM domains. **(a-c)** The full-length bMERB domain is required for MO-CH-LIM interaction. Binding affinities were measured by titrating the bMERB/bMERB_H1-2_/bMERB_H2-3_ domain (600 µM) with the MO-CH-LIM domain (40 µM). Integrated heat peaks, corrected for the offset by subtracting the control experiment, were fitted to a one-site-binding model, yielding the binding stoichiometry (N), enthalpy (ΔH), entropy (ΔS), and dissociation constant (K_D_). The data clearly show that the full-length bMERB construct is necessary for interaction with MO-CH-LIM. The data are representative of at least three repetitions. N.D. denotes not detected. (**d)** The Coomassie blue-stained SDS-PAGE analysis of purified Mical1 and the MO-CH-LIM:bMERB complex proteins, incubated in the absence or presence of the crosslinker disuccinimidyl dibutyric urea (DSBU), demonstrates efficient DSBU-dependent crosslinking. (**e-f)** Intramolecular crosslinking of Mical1. (**e)** Schematic visualization showing inter- and intra-domain crosslinks identified by mass spectrometry analysis of purified full-length Mical1 crosslinked by DSBU. Crosslinks are shown in light purple. (**f**) Schematic diagram showing inter and intra-domain crosslinks between purified MO-CH-LIM and bMERB domain crosslinked by DSBU. Schematic diagram illustrating inter- and intra-domain crosslinks between the purified MO-CH-LIM and bMERB domains, crosslinked by DSBU. Self-crosslinks are shown in light purple, whereas crosslinks between the isolated MO-CH-LIM and bMERB domains are shown in light blue.

To independently verify the interaction between the MO-CH-LIM and bMERB domains, we performed crosslinking mass spectrometry (XL-MS) analyses. Two experiments were conducted: first, using full-length Mical1; second, mixing the MO-CH-LIM construct and the bMERB domain in a 1:2 ratio to achieve a saturated MO-CH-LIM complex. (Fig. 2d-f). A total of 59 peptides were identified from Mical1 full-length experiments, and 76 peptides were identified from the MO-CH-LIM:bMERB complex. Crosslink mapping revealed numerous reproducible inter-domain and intra-domain connections between the two experiments. In the Mical1 full-length experiment, 6 peptides linked MO and bMERB domains, compared to 24 peptides in the MO-CH-LIM:bMERB complex experiment, confirming the interaction between the full-length bMERB domain and MO. This aligns with our ITC data, indicating the involvement of all three bMERB helices. However, no cross-links were found between CH and bMERB or LIM and bMERB domains. Our ITC results clearly demonstrate that the MO-CH-LIM construct interacts with the bMERB domain but not with MO or MO-CH. The absence of cross-links may be due to the limited number of lysine residues in the CH and LIM domains (2 in CH, 1 in LIM), potentially indicating they are not part of the binding interface. Nevertheless, XL-MS experiments suggest that the bMERB domain interacts with the MO domain, implying the formation of an intramolecular complex.

### Structural basis of Mical1 auto-inhibition

To understand the molecular basis of the MO-CH-LIM:bMERB interaction, we utilized AlphaFold2^31^ modeling to predict the auto-inhibited state of Mical1, producing a model with high confidence predicted local difference distance test (pLDDT) scores and predicted alignment errors (PAE), which we analyzed in detail (Fig. 3a and Supplementary Fig. 2).

**Fig. 3:**
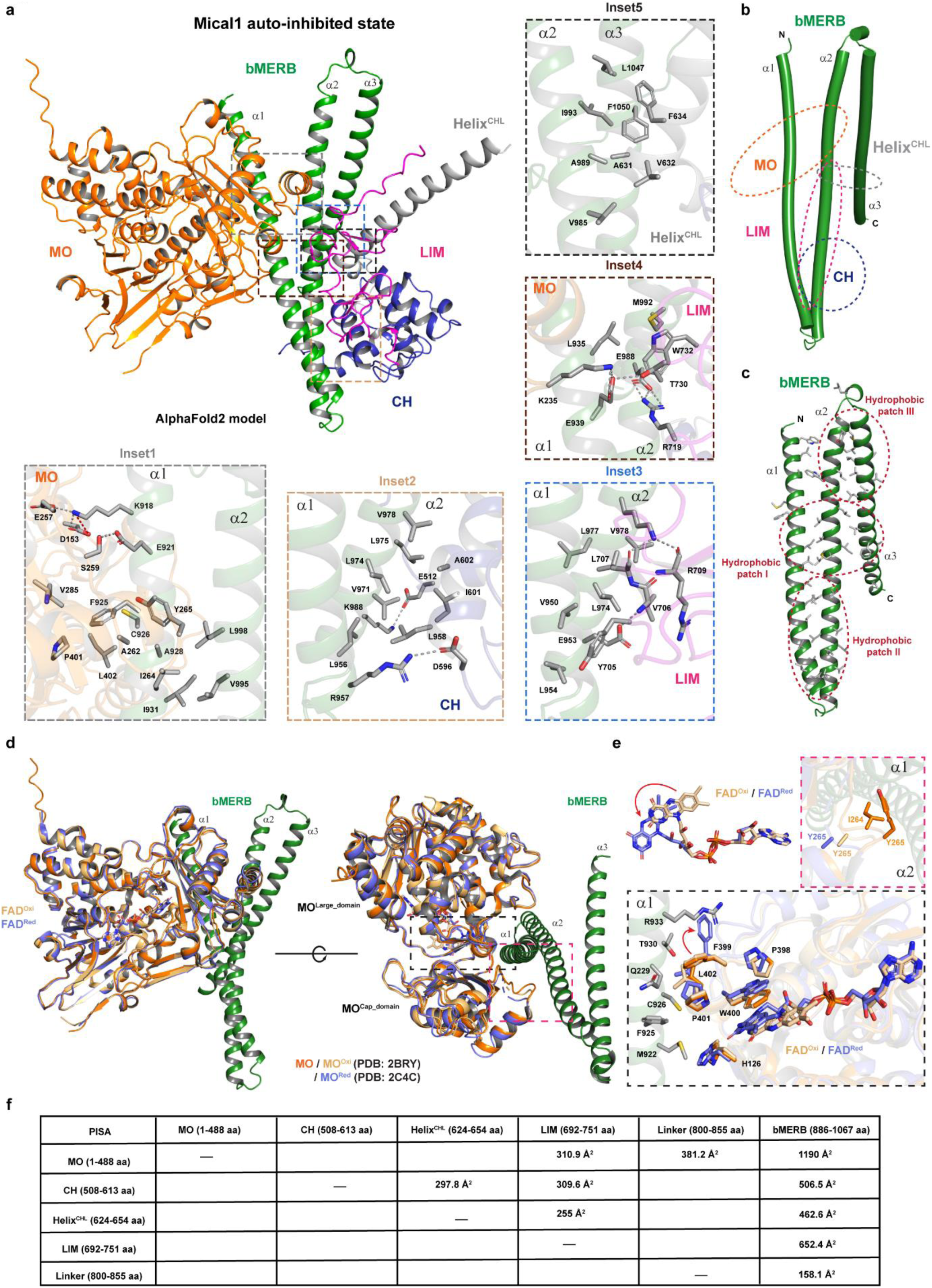
AlphaFold2 model of auto-inhibited Mical1. **(a)** The cartoon representation of the AlphaFold2 Mical1 model. The MO domain, CH domain, LIM domain, the helix connecting the CH and LIM domains (Helix^CHL^), and the bMERB domain of Mical1 are colored in orange, pink, blue, gray, and green, respectively. The insets highlight the various binding interfaces between the MO:bMERB, CH:bMERB, LIM:bMERB, and Helix^CHL^:bMERB domains, with hydrogen bonds and salt bridges indicated by gray dashed lines. (**b-c)** The cartoon representation of the bMERB domain depicts the MO domain, CH domain, LIM domain, and Helix^CHL^ binding interfaces in orange, pink, blue, and gray dashed lines, respectively. The hydrophobic patches are shown in red dashed lines on the bMERB domain. (**d)** Structural alignment was performed between the oxidized mouse Mical1 MO domain (light orange, PDB: 2BRY) and the reduced mouse Mical1 MO domain (slate, PDB: 2C4C), and both were compared with the human Mical1 AlphaFold2 model (MO domain in orange and bMERB domain in green). **(e)** Schematic illustration showing the movement of the isoalloxazine ring of the FAD head during the catalytic cycle upon NADPH treatment. Insets depict the conformational changes of several active site residues upon NADPH treatment. (**f)** Results from a systematic analysis of the binding interfaces between various domains of the AlphaFold2-predicted Mical1 auto-inhibited model using PISA^64^.

The AlphaFold2 model predicts that the MO (residues 1-488), CH (residues 508-613), Helix^CHL^ (residues 624-654), LIM (residues 692-751) domains, and linker (residues 800-855) interact with the C-terminal bMERB (residues 886-1067) domain, forming an auto-inhibited complex. Helices 1-2 of the bMERB domain interact with the MO, CH, and LIM domains, while helix 3 specifically interacts with Helix^CHL^, resulting in three distinct binding interfaces on the bMERB domain, predominantly through hydrophobic interactions (Fig. 3a-c). The first binding interface involves the first half of helix 1 and the second half of helix 2 of the bMERB domain, where the MO domain interacts (1190 Å^2^ buried surface). The MO domain consists of two core domains: a large domain (MO^Large_domain^, residues 1–226 and 374– 484) and a smaller cap domain (MO^Cap_domain^, residues 235–366). Both domains are connected by β-strands β9 (residues 228–234) and β15 (residues 367–373). The active site is a cleft formed by both the MO^Large_domain^ and the MO^Cap_domain^, containing a FAD molecule. Both the MO^Large_domain^ and the MO^Cap_domain^ interact with the bMERB domain. The MO^Large_domain^ interacts with helix 1, whereas the MO^Cap_domain^ interacts with both helices 1 and 2 (Fig. 3d,e). The binding site is opposite to the active site (FAD binding cleft). The second binding interface involves the center of the bMERB domain, where all three helices interact with the MO^Cap_domain^, LIM domain, and Helix^CHL^. The third binding interface centers on helices 1-2 of the bMERB domain, facilitating interactions between the CH (507 Å²) and LIM (652 Å²) domains (Fig. 3f). While helix 3 of the bMERB domain does not directly interact with the MO, CH, and LIM domains, our ITC data indicate its necessity for complex formation. This necessity is likely due to stabilizing interactions between helices 2 and 3 and the Helix^CHL^, suggesting that this interaction stabilizes the second binding interface where the MO and LIM domains interact with helices 1 and 2 of the bMERB domain (Fig. 3a).

Apart from the active site cleft, a smaller cleft is formed between the MO^Large_domain^ and MO^Cap_domain^, occupied by helix 1 of the bMERB domain (Fig. 3d). This cleft includes loops connecting β9-β10, β17-α15, and β15 of the MO^Large_domain^, and β11 and the loop connecting α10-11 of the MO^Cap_domain^. Helix 1 residue K918 forms hydrogen bonds with D153 and E257 of the MO domain, while E921 interacts with S259 of the MO domain (Fig. 3a, inset1). The side chain of F925 of the bMERB domain inserts itself into a hydrophobic patch formed by P401, L402, and V285 residues of the MO domain. Similarly, I264 of the MO^Cap_domain^ inserts into the hydrophobic patch involving helix 1 (I931) and helix 2 (V995 and L998), while Y265 of the MO^Cap_domain^ interacts hydrophobically with F925 (Fig. 3a, inset1). Additionally, K235 in the β9-β10 loop forms a hydrogen bond with E939 (Fig. 3a, inset4)

The CH and LIM domain binding sites on the bMERB domain span the C-terminal half of helix 1 and the N-terminal half of helix 2, characterized by a hydrophobic array. L598^CH^ interacts hydrophobically with L956, V971, and L974 of the bMERB domain, whereas I601^CH^ similarly engages with L975 and V978. Further, the CH:bMERB interaction is reinforced by hydrogen bonds between residues D596^CH^-R957 and E512^CH^-K988 (Fig. 3a, inset2). V632 and F634 of Helix^CHL^ interact hydrophobically with V985, I993, L1047, and F1050 of the bMERB domain (Fig. 3a, inset5). The LIM domain also engages extensively with the bMERB domain through hydrophobic interactions: Y705^LIM^ with V950 and L954, L707^LIM^ in the groove formed by L974, L977, and V978, and W732^LIM^ with L935 and M992. Additionally, hydrogen bonds are observed between R719^LIM^-E988 and T730^LIM^-E939 (Fig. 3a, inset3-4).

Next, we compared the auto-inhibited Mical1 model with previously published crystal structures of Mical1 individual domains^26,40–42^. The predicted MO domain closely resembles the isolated mouse Mical1 MO domain in both oxidized (PDB: 2BRY) and reduced states (PDB: 2C4C)^40^, with RMSD values of 0.299 Å for 396 Cα residues and 0.581 Å for 422 Cα residues, respectively (Supplementary Fig. 3a-b). Although the AlphaFold model lacks FAD, its active site residues are similar to the oxidized state of the mouse Mical1 MO domain (Fig. 3e). Even in the auto-inhibited state, the active site is accessible for catalysis. Previous studies have shown that in the reduced state of the MO domain, the FAD isoalloxazine ring flips during the catalytic cycle, leading to conformational changes in active site residues^40^. FAD’s isoalloxazine ring flips during the catalytic cycle in the reduced MO domain, causing changes in active site residues such as F399, W400, and P401 upon NADPH binding (Fig. 3d-e). F399, in particular, undergoes significant changes upon NADPH binding. In the auto-inhibited AlphaFold2 model, bMERB domain binding sterically hinders this rotation, stabilizing the oxidized MO domain and preventing the FAD transition to the reduced state (Fig. 3d-e). In the auto-inhibited state, the side chains of I264 and Y265 in the MO^Cap_domain^ are inserted into a hydrophobic patch formed by helices 1 and 2 of the bMERB domain. In the isolated oxidized/reduced mouse MO domain structure, Y265 and I264 lacked electron density, indicating that bMERB binding likely stabilizes the loop conformation of the MO domain (Fig. 3d-e).

Furthermore, structural analysis of isolated human CH (PDB: 2DK9) and bMERB domain (PDB: 5LE0) revealed differences from the AlphaFold2 predictions, with RMSD values of 1.089 Å for 92 Cα residues in the CH domain and 0.967 Å for 94 Cα residues in the bMERB domain^26,41^. Notably, the maximum structural difference was observed for the isolated human LIM domain (PDB: 2CO8), with an rmsd of 2.135 Å for 65 Cα residues^42^ (Supplementary Fig. 3c-f). We hypothesize that the bMERB domain first interacts with the CH domain, establishing the binding interface for the LIM domain. As the LIM domain binds, it brings all three domains together, stabilizing the MO binding interface and forming a compact structure. Finally, the MO domain interacts with both the bMERB and LIM domains, facilitating the formation of a tightly packed auto-inhibited complex.

### Generation of bMERB interface mutants

To corroborate the Mical1 auto-inhibited AlphaFold2 model, we conducted biochemical experiments. Sequence alignment of bMERB domains reveals that surface residues interacting with MO-CH-LIM are highly conserved across mammalian homologs (Supplementary Fig. 4). To assess the significance of individual interface residues, we introduced mutations into each helix of the bMERB domain based on the AlphaFold2 Mical1 auto-inhibited model and analyzed their interaction with MO-CH-LIM using ITC experiments. (Fig. 4). All bMERB mutants with alanine substitutions at conserved residues, except K918A_E921A, show a significant decrease in binding affinity. The K918A and E921A double mutations in helix 1 reduce affinity for MO-CH-LIM by only 2-fold (Fig. 4c). K918 forms hydrogen bonds with D153 and E257 of the MO domain, while E921 bonds with S259. In contrast, mutation of F925 abrogates the MO-CH-LIM interaction (Fig. 4d), as its side chain inserts into the hydrophobic pocket formed by A262, Y265, and V285 of the MO domain. The E939A mutant shows more than a 5-fold reduction in binding affinity with MO-CH-LIM (Fig. 4e). The side chain of E939 forms hydrogen bonds with K235 of the MO domain and T730 in the LIM domain.

**Fig. 4:**
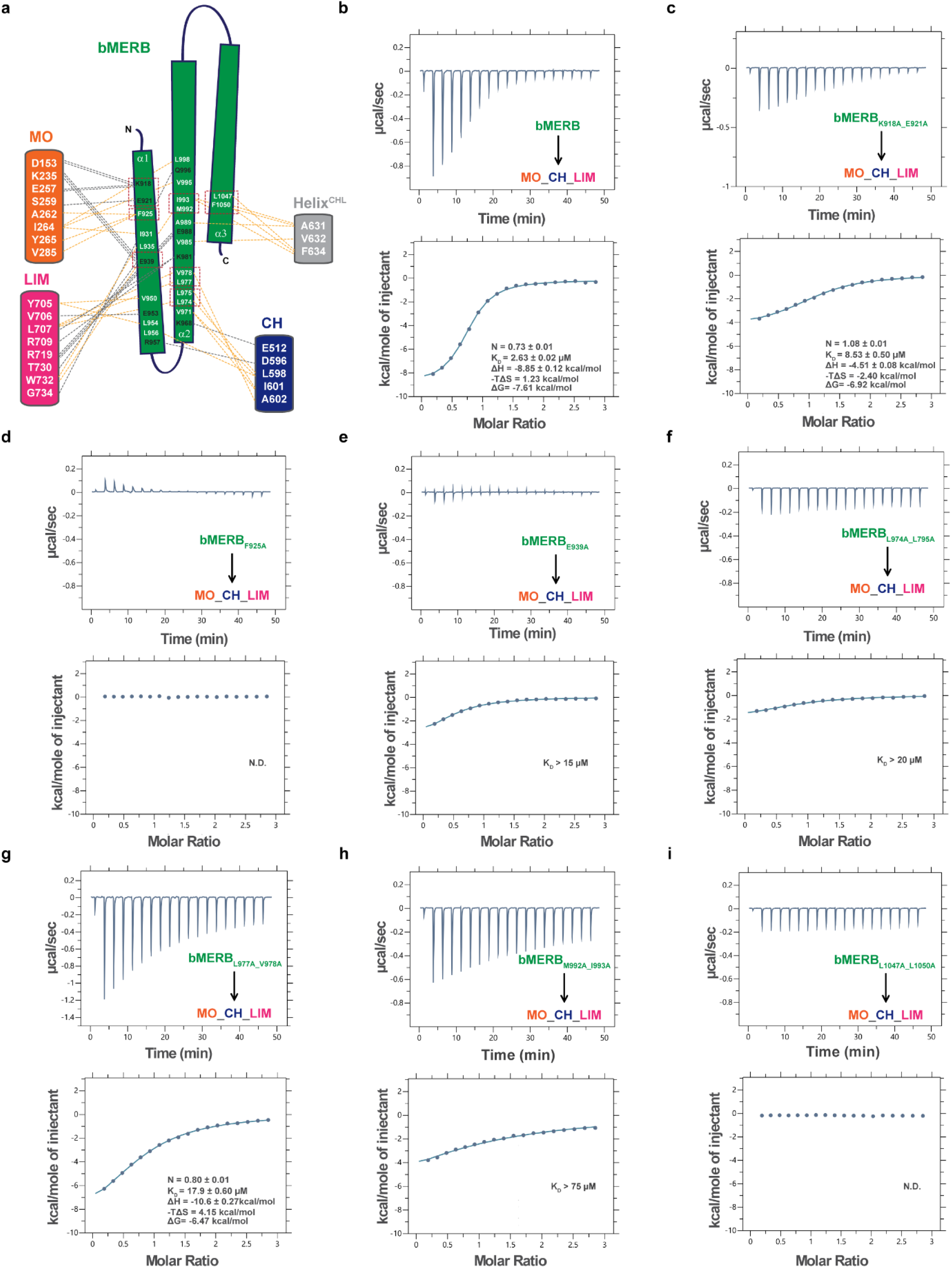
Crucial bMERB interface residues for MO-CH-LIM interaction. **(a)** Schematic illustration of the interactions between the MO-CH-LIM and the bMERB domain. Hydrophobic interactions are represented by light orange dashed lines, while hydrogen bonds (H-bonds) and salt bridges are depicted with gray dashed lines. Residues that have been mutated are highlighted in red dashed boxes. (**b**–**i)** Mutational alanine screening of the bMERB domain via ITC measurements. The binding of different bMERB mutants (600 µM) with MO-CH-LIM (40 µM) was systematically tested and affinities were determined by ITC experiments. Integrated heat peaks, corrected for the offset by subtracting the control experiment, were fitted to a one-site-binding model, yielding the binding stoichiometry (N), enthalpy (ΔH), entropy (ΔS), and dissociation constant (K_D_). N.D. denotes not detected. No binding was detected for the mutant F925A or the mutant L1047A_F1050A. However, a more than 5-fold reduction in binding affinity was observed for all other mutants, except for K918A_E921A, which exhibited a marginal defect. The data represent at least three repetitions.

Double alanine mutations in helix 2 (L974A_L975A and L977A_V978A) lead to a 6-fold reduction in binding affinity (Fig. 4f, g). Residues L956, V971, L974, L975, L977, and V978 of the bMERB domain form a hydrophobic patch that accommodates the CH and LIM domains. The side chain of L598 in the CH domain fits into the hydrophobic pocket formed by L956, V971, and L974, while I601 interacts with V971 and L975. Additionally, L707 of the LIM domain forms a hydrophobic interaction with L974, L977, and V978 of the bMERB domain. The M992A_I993A double alanine mutant in helix 2 causes a more than 25-fold reduction in binding affinity (Fig. 4h). M992 interacts with LIM domain residue W732, and I993 interacts with F634 of Helix^CHL^ (residues 624-654).

Furthermore, mutating residues L1047 and F1050 of helix 3 to alanine eliminates detectable binding with MO-CH-LIM (Fig. 4i). Notably, L1047 and F1050 interact with Helix^CHL^ (residues 624-654), not with MO, CH, or LIM domains. The side chain of F634 of Helix^CHL^ inserts into a hydrophobic patch formed by I993 (helix 1), L1047, and F1050 (helix 2) of the bMERB domain, while V632 of Helix^CHL^ interacts with V985 of bMERB. The L1047 and F1050 mutations destabilize this hydrophobic patch.

### Allostery between the two Rab binding site

We previously demonstrated that the Mical1 bMERB domain binds Rab8 in a 1:2 stoichiometry, with helices 1-2 forming the low-affinity N-terminal binding site (Nt-RBS1) and helices 2-3 forming the high-affinity C-terminal binding site (Ct-RBS2)^28^. Deleting either the N- or C-terminal helix resulted in binding Rab molecules at a 1:1 stoichiometry^28^. We investigated whether two Rab molecules bind to the bMERB domain of full-length Mical1 and as expected full-length Mical1 binds Rab8 with a 1:2 stoichiometry, similar to the isolated bMERB domain, but with slightly lower affinity (K_D1_: 40 vs. 10 nM and K_D2_: 0.73 vs. 1.12 µM) (Fig. 5a-b). Next, we examined whether the binding of a Rab molecule at either the N- or C-terminal site affects the binding of a second Rab molecule at the other site, or if the two sites operate independently. Since full-length Mical1 exhibits a similar range of Rab8 binding affinities as the isolated bMERB domain (Fig. 5b, h), we generated and purified isolated bMERB domain mutants based on the bMERB:Rab10 complex (PDB: 5LPN)^28^ structure (Fig. 5c). We assessed their interaction with Rab8 by varying the bMERB:Rab ratio (1:1.1 and 1:2.2) using analytical size exclusion chromatography. These experiments aimed to identify constructs that either show no or defective complex formation or fail to display an increase in complex formation upon increasing the Rab concentration. Furthermore, the stoichiometry of the complex was determined using isothermal titration calorimetry (ITC) with selective constructs.

**Fig. 5:**
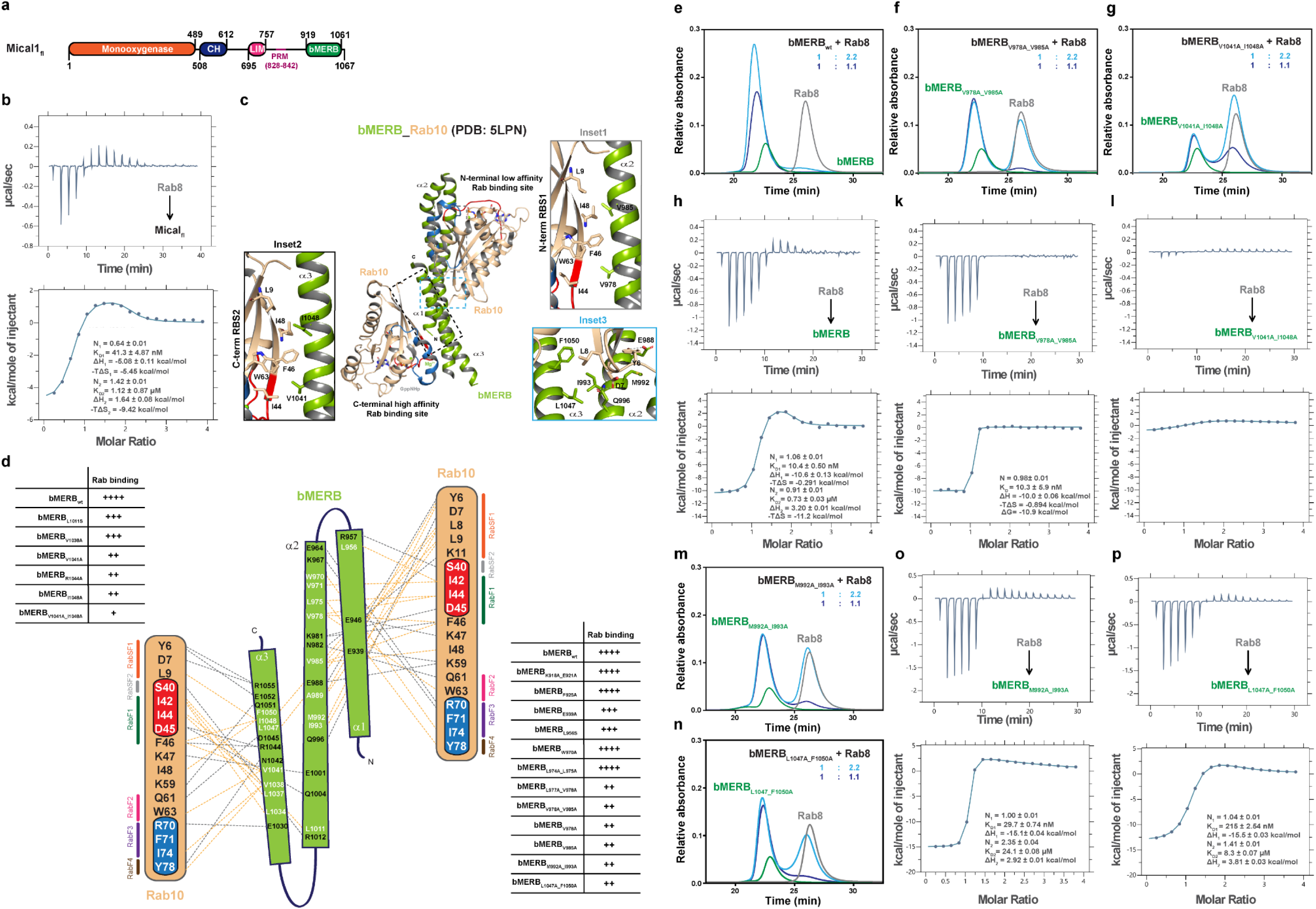
Allostery between the Rab binding domains. **(a)** A schematic diagram of human Mical1. **(b)** Full length Mical1 binds to Rab8 in 1:2 stoichiometry. Binding affinities were measured by titrating the Rab8 (800 µM) to full-length Mical1 (40 µM). Integrated heat peaks were fitted to a two-site-binding model yielding the binding stoichiometry (N), the enthalpy (ΔH), the entropy (ΔS), and the dissociation constant (*K*_D_). **(c)** Cartoon representation of the bMERB:Rab10 complex structure (PDB: 5LPN)^28^, with the bMERB domain colored in green and the Rab10 domain in wheat. **(c-d)** Schematic illustration of the interactions between the bMERB domain and Rab10. Rab10 binds to the Mical1 bMERB domain via its N-terminal regions, switch regions, and inter-switch region. Hydrogen bonds and salt bridges are depicted with gray dashed lines, while light orange dashed lines indicate hydrophobic interactions. RabSF1, RabSF2, RabF1, RabF2, RabF3, and RabF4 motifs are colored in orange, gray, green, pink, purple, and brown, respectively. Additionally, analytical size exclusion chromatography results are provided from a systematic analysis of interactions between the bMERB domain mutants and increasing concentrations of Rab8 (refer to Supplementary Fig. 4 and 5). (**e-g, m-n)** The bMERB domain (green), Rab8 (gray), and a mixture of both (blue) were loaded onto a Superdex 75 10/300 GL column to monitor for complex formation. (**h-l, o-p)** Binding affinities were measured by titrating the Rab8 (600 µM) to mutants bMERB domain (30 µM). Integrated heat peaks were fitted to a one-site-binding or two-site-binding model yielding the binding stoichiometry (N), the enthalpy (ΔH), the entropy (ΔS), and the dissociation constant (*K*_D_). The data are representative of at least three repetitions.

First, we generated mutants targeting Nt-RBS1. Mutants K918A_E921A, F925A, W970A, and L974A_L975A showed no binding defects (Supplementary Fig. 5a-d, h). However, E939A and L956S exhibited only a slight increase in complex formation at higher bMERB ratios (1:1.1 to 1:2.2), indicating a lower affinity for the second N-terminal Rab binding (Supplementary Fig. 5e, f). Mutants L977A_V978A, V978A_V985A, V978A, V985A, M992A_I993A, and L1047A_F1050A showed no significant change in complex formation with increased Rab concentration, suggesting either 1:1 binding or defective binding at both sites (Supplementary Fig. 5i-n).

We next generated mutants targeting Ct-RBS2. Mutants L1011S and V1038A showed no apparent defects (Supplementary Fig. 6a-d). However, mutants V1041A, R1044A, and I1048A formed smaller complexes on aSEC and showed no significant change in complex formation with varying Rab concentrations, suggesting binding defects (Supplementary Fig. 6e-g). Notably, the double mutant V1041A_I1048A showed minimal complex formation, indicating that these mutations affect binding at both Rab sites (Supplementary Fig. 6h).

Distinct heat exchange patterns for Rab binding at the N-terminal and C-terminal sites of Mical1 were previously observed, with C-terminal binding being enthalpy-driven and N-terminal binding entropy-driven^28^. To precisely determine the impact of specific mutations on Rab binding, we conducted ITC experiments with mutants: bMERB_V978A_V985A_, bMERB_V1041A_I1048A_, bMERB_M992A_I993A_, and bMERB_L1047A_F1050A_. The ITC data unequivocally demonstrate that bMERB_V978A_V985A_ binds to Rab8 with a 1:1 stoichiometry, an enthalpy-driven reaction, and affinity similar to Ct-RBS2. In contrast, bMERB_V1041A_I1048A_ exhibited low heat exchange and cannot be reliably fitted, indicating that C-terminal binding is necessary for Rab binding at the N-terminal site (Fig. 5h-l). Residues V978/V985 and V1041/I1048 are functionally equivalent, interacting with specific Rab10 residues (I44, W63, F46 for V978/V1041; L9, F46A, I48 for V985/I1048) (Fig. 5c, Insets 1-2). V978 and V985 are crucial for N-terminal Rab binding, as single mutants V978A and V985A show similar binding as of the double mutant V978A_V985A. Conversely, the double mutant V1041A_I1048A exhibits a pronounced effect on Rab10 binding compared to individual mutants V1041A and I1048A, indicating that perturbations in the N-terminal binding site still allow Rab8 to bind to the C-terminal site, whereas perturbations at Ct-RBS2 also affect binding at Nt-RBS1. This suggests a sequential binding mechanism for Rabs. To further support our observation, we performed SEC experiments with V978A_V985A and V1041A_I1048A bMERB mutants with other Rab8 family members (Rab10, Rab13, and Rab15). Similar observations were made; however, compared to Rab8, Rab10 showed a lesser increase in complex formation upon increasing the bMERB:Rab ratio, suggesting a slightly lower affinity for the N-terminal Rab binding (Supplementary Fig. 7).

The other Nt-RBS1 mutant, bMERB_M992A_I993A_, exhibited a similar high affinity for C-terminal Rab8 binding, while a more than 30-fold decrease in N-terminal Rab binding was observed. The residue M992 forms hydrophobic interactions with Y6 of Rab10 (Nt-RBS1) (Fig. 5o, Inset3). An intriguing observation arose with the mutant bMERB_L1047A_F1050A_, which demonstrated reduced binding affinity for both Rab binding sites. This mutant showed over a 20-fold decrease in C-terminal Rab8 binding and a 10-fold reduction in N-terminal Rab binding (Fig. 5p). Residues L1047 and F1050 contribute to hydrophobic interactions with L8 of Rab10 (Nt-RBS1) (Fig. 5c, Inset3). However, L1047 and F1050 do not participate in binding with Rab10 at Ct-RBS2; the decrease in binding affinity might be due to a change in local hydrophobicity, as these residues lie on the opposite side of the crucial I1048. Previously, we reported that the N-terminus of the Rab8 family member provides specificity^28^. L8^Rab10^ is implicated in the RabSF1 interaction with L1047 and F1050, thereby further stabilizing the interaction. These findings suggest that the conformation of helix 3 of the bMERB domain plays a crucial role in Rab binding at both the N-terminal and C-terminal Rab binding sites within the full-length bMERB domain background. This contrasts with earlier findings showing that Rab8 bound to N- and C-terminal helix-deleted constructs in a 1:1 ratio, suggesting site independence^28^.

We propose that the absence of helix 3 may cause helices 1-2 to adopt a different conformation, accommodating the Rab molecule. Overall, our data suggest allosteric communication between the N-terminal and C-terminal Rab binding sites, with the first Rab molecule binding at Ct-RBS2, followed by the second Rab molecule binding at Nt-RBS1.

### The overall structure of the bMERB_V978A_V985_:Rab10/bMERB_V978A_:Rab10 complex

To gain mechanistic insight, we attempted to crystallize bMERB_V978A_V985_:Rab10, bMERB_V978A_:Rab10, bMERB_V985A_:Rab10 complexes, and the bMERB_V1041A_I1048A_ domain. We successfully obtained well-diffracting crystals of the bMERB_V978A_V985_:Rab10 and bMERB_V978A_:Rab10 complexes, with space group P2_1_, diffracting to 1.8 and 2.05 Å, respectively. Structures were solved as described in the materials and methods (Data and refinement statistics are in Supplementary Table 1).

The asymmetric unit contained a single copy of either the bMERB_V978A_V985A_:Rab10 or bMERB_V978A_:Rab10 complex. Both structures are nearly identical (Fig. 6a and Supplementary Fig. 8), so here we describe the bMERB_V978A_V985A_:Rab10 complex structure in detail. In accordance with our aSEC and ITC data, a single molecule of Rab10 was bound to the C-terminal Rab binding site. Structural alignment of bMERB_V978A_V985A_:Rab10 with the wild-type bMERB:Rab10 (PDB: 5LPN)^28^ shows that the C-terminal Rab binding site of the mutant bMERB domain adopts the same conformation as the wild-type complex. However, the second half of helix 2 of the mutant bMERB domain adopts a different conformation. The N-terminal Rab binding site is formed by a kink in the second half of helix 2 and the movement of the first half of helix 1 towards Rab. In the bMERB_V978A_V985A_:Rab10 structure, the second helix’s conformation is similar to the unbound bMERB domain conformation (PDB: 5LEO)^26^ (Fig. 6b-c). This mutation of V978 or V985 leads to the abrogation of the second Rab binding at the N-terminal Rab binding site, as found in aSEC/ITC experiments as well.

**Fig. 6:**
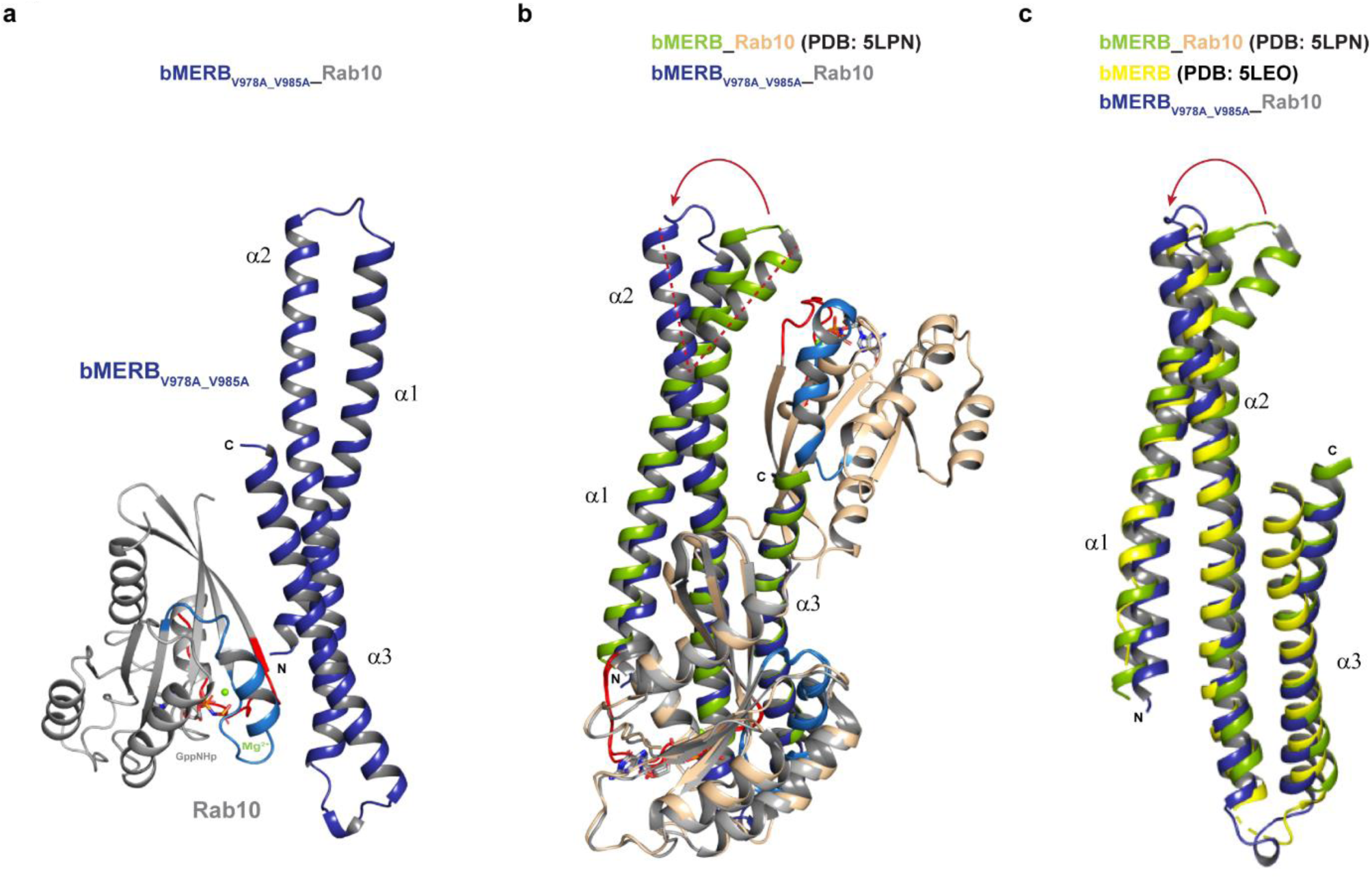
Crystal structure of the bMERB_V978A_V985A_:Rab10 complex. **(a)** Cartoon depiction of the bMERB_V978A_V985A_:Rab10 complex. A single Rab10 molecule (gray) binds to the bMERB_V978A_V985A_ domain (blue) at the high-affinity C-terminal Rab-binding site. Switch I and Switch II of Rab10 are shown in red and blue, respectively. GppNHp and Mg^2+^ are depicted as sticks and a green sphere, respectively. **(b)** Structural superposition of the bMERB_V978A_V985A_:Rab10 complex and bMERB:Rab10 complex (PDB: 5LPN)^28^. **(c)** Structural alignment of the bMERB domains of bMERB_V978A_V985A_:Rab10 (blue) with the bMERB:Rab10 (PDB: 5LPN)^28^ and the bMERB domain (PDB: 5LEO)^26^.

### Phosphomimetic mutation of S960 of the bMERB domain

Recently, Macgarry *et al*., proposed an alternate mode of Mical1 activation by Rho GTPases and showed that upon extracellular ligand stimulation, Cdc42 interacts with its effector PAK1 and active Cdc42:PAK1 then binds to the MO domain of Mical1, PAK1, a serine/threonine kinase, phosphorylates S817 and S960 of the Mical1 bMERB domain^43^. The authors proposed that this phosphorylation enhances Rab10 binding, which activates Mical1 and increases F-actin disassembly^43^. We analyzed AlphaFold2 model and found that S817 lies in the linker region, while S960 is part of the bMERB domain. Despite the low pLDDT score for this linker region, the position of S817 is highly reproducible across models, and the side chain is pointing towards a negatively charged patch. Phosphorylation might induce charge repulsion, potentially ’loosening’ the auto-inhibited complex (Fig. 7a-b and Supplementary Fig. 9). However, further biochemical and structural validation is needed.

**Fig. 7:**
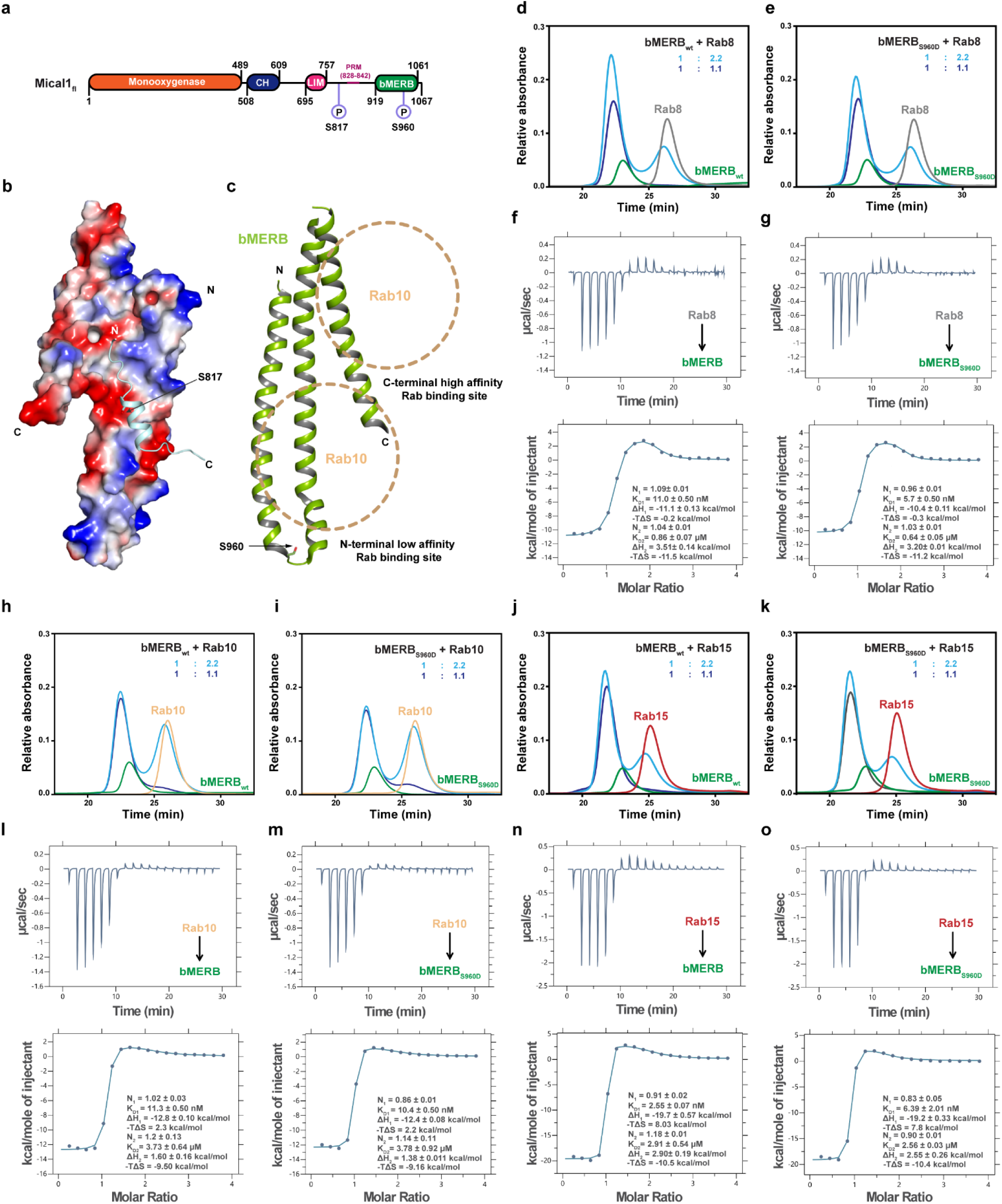
Phosphomimetic mutation of bMERB S960D does not affect Rab8 family member binding. **(a)** The schematic diagram of human Mical1 shows an N-terminal monooxygenase (MO) domain, a calponin homology (CH) domain, a Lin11, Isl-1, and Mec-3 (LIM) domain, and a C-terminal coiled-coil bMERB (bivalent Mical/EHBP Rab binding) domain, along with PAK1 phosphorylation sites S817 and S960. **(b)** The surface electrostatic potential of bMERB was calculated in PyMOL using the APBS-PDB2PQR plugin and visualized with colors ranging from red (-5 kT/e) to blue (+5 kT/e). AlphaFold2 predicted the Mical1 amino acids 807–820, with S817 shown in stick representation. The side chain of S817 points towards a negatively charged patch on the Mical1 bMERB domain. **(c)** Cartoon representation of the Mical1 bMERB domain (PDB: 5LPN)^28^, with S960 shown in stick representation. Rab10 binding sites are indicated with dashed circles. **(d, e, h-k)** Binding of Rab8a (gray)/Rab10 (wheat)/Rab15 (red) with different Mical1 bMERB domain (green) was systematically tested onto a Superdex 75 10/300 GL column. **(f, g, l-o)** Binding affinities were measured by ITC experiments. Binding affinities were measured by titrating the Rab8/Rab10/Rab15 (600 µM) to wild type/mutant bMERB domain (30 µM). Integrated heat peaks were fitted to a two-site-binding model yielding the binding stoichiometry (N), the enthalpy (ΔH), the entropy (ΔS), and the dissociation constant (*K*_D_). Phosphomimetic mutation of S960 of the bMERB domain does not affect interaction with Rab8/Rab10/Rab15. The data are representative of at least three repetitions.

Previously, we showed that the bMERB domain interacts with Rab8, Rab10, Rab13, and Rab15, but lacked quantitative binding data. We have now quantified these interactions. Similar to Rab8, ITC data reveal that Rab10 and Rab15 bind to the bMERB domain in a 1:2 stoichiometry, with one high-affinity site (Ct-RBS2: 11 nM for Rab10, 2.5 nM for Rab15; enthalpy-driven) and one low-affinity site (Nt-RBS1: 3.73 µM for Rab10, 2.91 µM for Rab15; entropy-driven). We purified the bMERB_S960D_ mutant and assessed its interaction with Rab8, Rab10, and Rab15 using aSEC/ITC. The phosphomimetic mutation at S960 did not affect Rab binding, consistent with our structural observations that S960, located in helix 1 of the Nt-RBS1 site, does not interact with Rab10 (Fig. 7c-o).

Based on our findings, we propose that phosphorylation of both S817 and S960 may facilitate the release of auto-inhibition in Mical1, exposing the bMERB domain for Rab binding without affecting Rab affinity. Further biochemical studies are needed to fully elucidate the activation mechanism of Mical1 by the RhoGTPase-PAK1 pathway.

### Impact of Rab binding site mutations in the bMERB domain on MO-CH-LIM interaction

In the absence of Rab, the Nt-RBS1 of bMERB accommodates the CH and LIM domain, so we investigated the effect of several RBS mutants of the bMERB domain on MO-CH-LIM interaction (Fig. 8). The L956S mutant displayed more than a 9-fold reduction in binding affinity (Fig. 8c). L956 forms a hydrophobic interaction with L598 of the CH domain; however, it is also part of a continuous array of hydrophobic residues. Therefore, we propose that this mutation may lead to the destabilization of the hydrophobic patch. In contrast, the phosphomimetic mutant S960D did not affect MO-CH-LIM binding, as this residue is not part of the predicted binding interface (Fig. 8d). Another putatively non-interacting residue, the W970A mutant, exhibited a 4-fold decrease in binding affinity (Fig. 8e). Although W970^bMERB^ is not directly involved in the interaction, it may contribute to stabilizing the hydrophobic patch of the bMERB domain, which is crucial for MO-CH-LIM interaction.

**Fig. 8:**
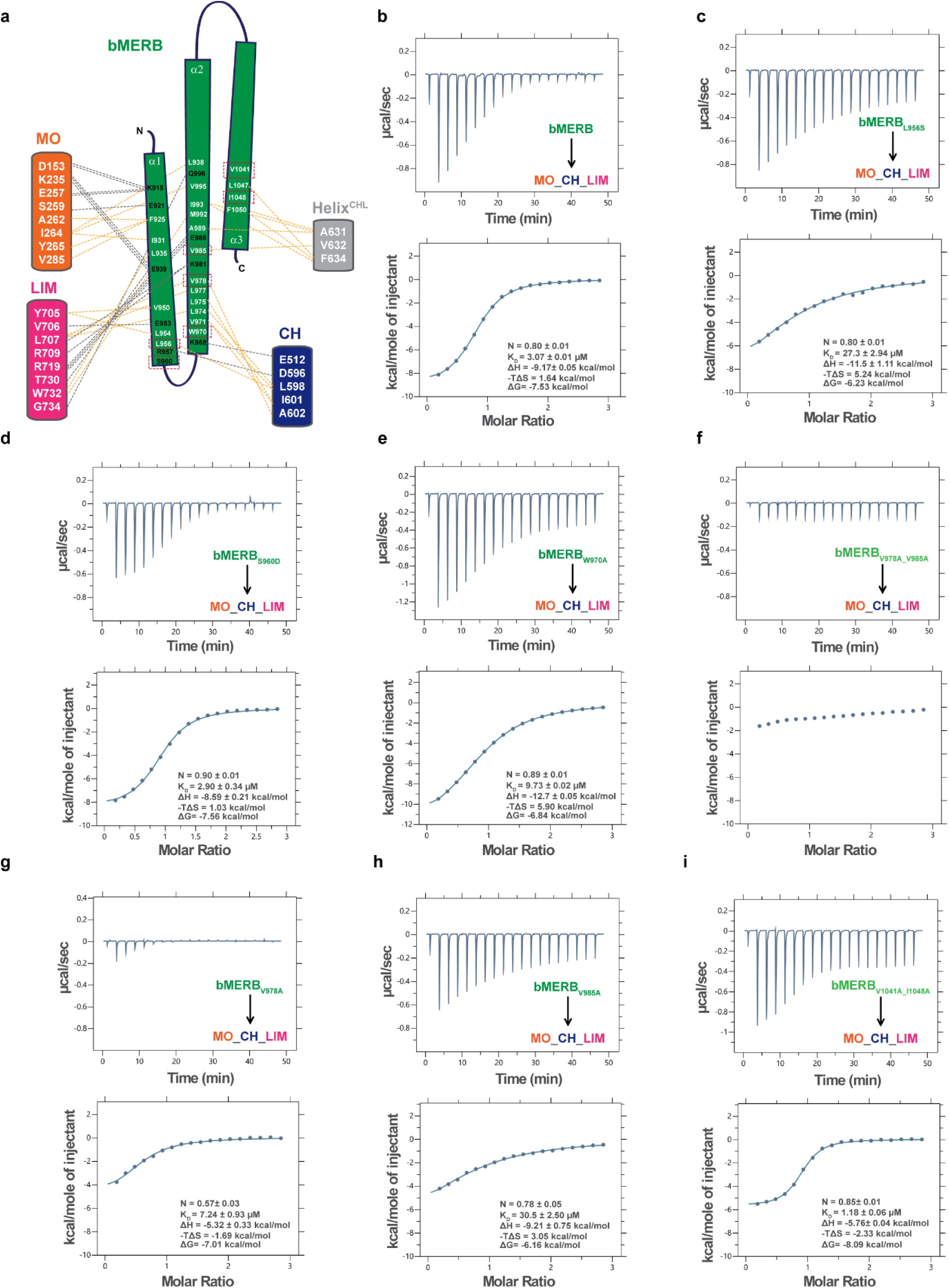
Impact of Rab binding site mutations in the bMERB domain on MO-CH-LIM interaction. **(a)** Schematic illustration of the interactions between the MO-CH-LIM domain and the bMERB domain. Hydrophobic interactions are indicated by light orange dashed lines, and hydrogen bonds are indicated by gray dashed lines. **(b-i)** Effect of RBS mutants on MO-CH-LIM:bMERB domain interaction. The binding of MO-CH-LIM with different RBS bMERB mutants was systematically tested, and affinities were determined by ITC experiments. Integrated heat peaks, corrected for the offset by subtracting the control experiment, were fitted to a one-site-binding model, yielding the binding stoichiometry (N), enthalpy (ΔH), entropy (ΔS), and dissociation constant (K_D_). N.D. denotes not detected. The data are representative of at least three repetitions.

The double alanine mutations V978A_V985A completely abolished the MO-CH-LIM interaction (Fig. 8f). Interestingly, single mutations V978A and V985A caused a 2-fold and 10-fold reduction in binding affinity, respectively (Fig. 8g-h). V978 interacts with A602 of the CH domain and L707 of the LIM domain, while V985 interacts with V632 of the Helix^CHL^. This indicates that the hydrophobic nature of these residues is crucial for stabilizing CH and LIM interactions with the bMERB domain. In contrast, double alanine mutations of non-interface residues, V1041A_I1048A, showed slightly better affinity than the wild type (Fig. 8i).

### Structural basis of Mical1 activation

To unravel the structural basis for the release of the MO, CH, and LIM domains from the bMERB domain upon Rab binding, we superimposed the Mical1 auto-inhibited AlphaFold2 model with the bMERB:Rab10 complex (PDB: 5LPN) structure^28^. Superimposition reveals that in the auto-inhibited Mical1 model, the Nt-RBS1 site is blocked by the CH and LIM domains, along with Helix^CHL^. Upon Rab binding, the bMERB domain helices undergo significant conformational changes (Fig. 9a-c). Helix 3 adopts a new conformation that interferes with the CH domain and Helix^CHL^. Meanwhile, binding of the second Rab molecule shifts helices 1-2 towards the Rab, further disrupting the interaction with the CH domain.

**Fig. 9:**
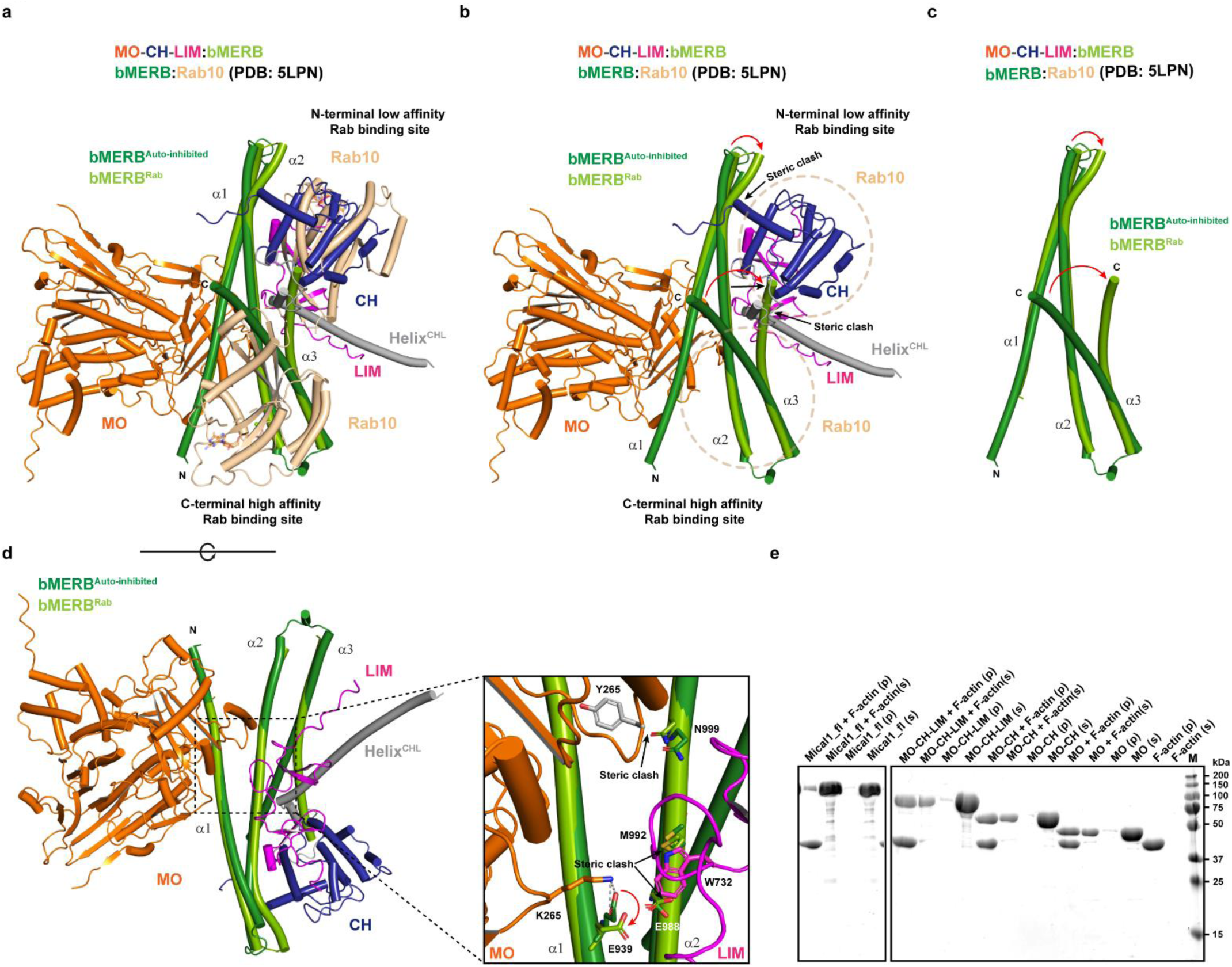
Structural basis of Mical1 activation upon Rab8 binding. **(a)** Structural superposition of the Mical1 auto-inhibited AlphaFold2 model and the bMERB:Rab10 complex (PDB: 5LPN)^28^. (**b)** Structural overlay clearly shows that the binding of the first Rab molecule at high affinity stabilizes the helix 3 conformation, which structurally interferes with the CH domain and Helix^CHL^ binding site. Additionally, the movement of helices 1-2 strictly hinders the CH binding site. **(c)** The superimposition of the bMERB domain of Mical1 in the auto-inhibited AlphaFold2 model with the bMERB domain of the bMERB:Rab10 complex (PDB: 5LPN)^28^ reveals the movement of bMERB helices upon Rab binding. **(d)** Structural overlay of the bMERB domain of Mical1 in the auto-inhibited state Alphafold2 model with the bMERB domain of the bMERB:Rab10 complex (PDB: 5LPN)^28^, with an inset highlighting the conformational changes of interface residues upon Rab10 binding. **(e)** The results of systematic analysis of interactions between full-length Mical1 or different Mical1 deletion constructs with F-actin via co-sedimentation experiments reveal that the MO domain is the minimal domain required for interaction with F-actin, whereas the full-length Mical1 exhibits perturbed interaction with F-actin. These experiments were independently repeated at least three times with consistent results.

The AlphaFold2 auto-inhibited model suggests that while the active site is accessible, helix 1 of the bMERB domain restricts the catalytic loop (amino acids 395-405) movement (Fig. 3e). Conformational changes in W400 stabilize FAD binding via π-π stacking and may facilitate hydride transfer from NADPH to FAD^40^. Several studies suggest that during oxidation, methionine residues approach W400, making them accessible to hydrogen peroxide^40,44^. The AlphaFold2 model indicates that enzyme activity is likely inhibited by the bMERB domain, which restricts the movement of the catalytic loop (Fig. 3e). In the Mical1 auto-inhibited model, F399’s movement is restricted, and the MO domain conformation resembles that of the isolated MO domain in the oxidized state (PDB: 2BRY) (Fig. 3e).

Next, we propose a model for Mical1 activation (Fig. 10). Comparing the Mical1 auto-inhibited state with the bMERB complex reveals that Nt-RBS1 is blocked by the CH, Helix^CHL^, and LIM domains. However, helix 3 of the bMERB domain is accessible, allowing the first Rab8 molecule to bind at the C-terminal high-affinity Rab binding site, defining the helix 3 conformation. This binding displaces the CH domain and Helix^CHL^, loosening the structure and exposing Nt-RBS1. A second Rab molecule then binds at the low-affinity RBS, preventing the reassembly of the bMERB:CH domain. The binding induces significant conformational changes in helices 1 and 2, disrupting their interaction with the MO domain. Key residues E939, F925, and N999 in helices 1 and 2 shift side chain orientations, leading to MO domain dissociation (Fig. 9d), supported by ITC data. These movements also further dissociate the LIM domain. Consequently, the binding of the second Rab molecule relieves auto-inhibition, allowing access to the MO, CH, and LIM domains. The active MO domain then induces F-actin depolymerization.

**Fig. 10:**
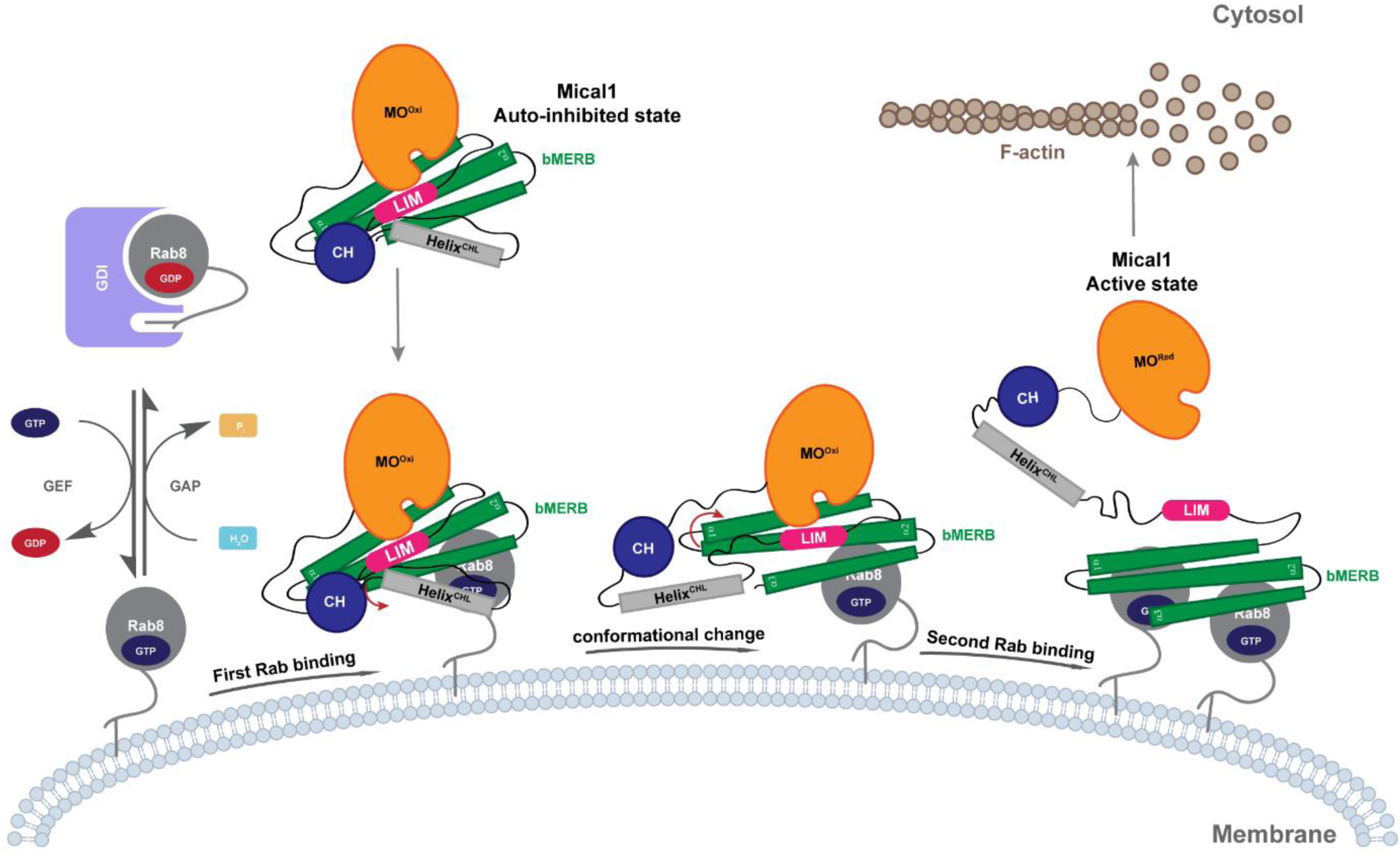
The activation mechanism of Mical1 by Rab8 family members. In the absence of active Rab8, Mical1 exists in an auto-inhibited state, where the MO, CH, Helix^CHL^, and LIM domains interact with its bMERB domain. Rab8 is recruited to the membrane by its GEF molecule (e.g., Rabin 8 or GRAB). The first molecule of active Rab8 binds to the high-affinity C-terminal binding site, leading to a change in the conformation of helix 3, which displaces the CH domain and Helix^CHL^. The first binding of the Rab molecule also changes the conformation of helix 2 and helix 3 at the low-affinity N-terminal Rab binding site, further interfering with CH binding, and this conformation leads to a change in the LIM and MO binding interface on the bMERB domain, resulting in the displacement of the LIM and MO domains. The second Rab8 molecule binds to stabilize the active state of the enzyme, allowing the MO domain to be available for F-actin interaction, leading to the depolymerization of F-actin.

Mical1 regulates actin dynamics by selectively oxidizing Met44 and Met47 of F-actin in an NADPH-dependent manner, causing depolymerization and severing of F-actin^6,45,46,49^. We have previously shown that Mical1’s CH, LIM, or CH-LIM domain fails to interact with F-actin ^30^. The Mical1 CH domain is structurally similar to CH type-2 domains, which cannot directly bind to F-actin, unlike CH type-1 domains. We tested full-length Mical1, MO-CH-LIM, MO-CH, and MO constructs for F-actin interaction using actin co-sedimentation assays. The results showed that MO-CH-LIM, MO-CH, and MO constructs interact with F-actin, indicating the MO domain is essential and sufficient for this interaction. Full-length Mical1 showed less interaction, suggesting that Rab binding is needed to overcome auto-inhibition for F-actin binding (Fig. 9e).

Previously, we suggested that the binding of two Rab molecules to Mical1 could increase the local concentration of specific Rab molecules, creating Rab microdomains and controlling Mical1 activity. Based on our data, we now propose that the initial binding of the first Rab molecule to a high-affinity site recruits Mical1 to specific membranes and alleviates its auto-inhibited state. This facilitates the subsequent binding of the second Rab molecule at the low-affinity Nt-RBS1 site, further stabilizing Mical1’s active state by preventing the rebinding of the MO, CH, and LIM domains. This process regulates the duration of enzyme activity and thus F-actin dynamics (Fig. 10). For Mical-like proteins, a concerted Rab recruitment cascade has been reported. For instance, Mical-L1 connects Rab35 and Rab8. Another study demonstrated that Mical-L1 is recruited to recycling endosomes by Rab35, which subsequently recruits other Rab proteins such as Rab8, Rab13, and Rab36^50^. Similarly, in Mical1-L2-dependent GLUT4 translocation, trafficking relies on the concerted action of Rab8 and Rab13^51^. The authors suggested that bMERB domain dimerization allows for concerted recruitment; however, in the case of Mical1, a single bMERB domain can recruit Rab proteins in a concerted manner.

In conclusion, our study has provided biochemical and structural insights into both the auto-inhibited and active states of Mical1. Our data offer a detailed overview of the MO-CH-LIM:bMERB interaction, with mutagenesis experiments highlighting the crucial residues involved. Additionally, we have elucidated the activation mechanism of Mical1, presenting evidence of allosteric regulation between the two Rab binding sites and identifying the key residues required for Rab recruitment.

## Methods

### Plasmid cloning

All expression constructs were generated by standard cloning techniques, using Phusion polymerase, restriction digestion, and ligation by T4 DNA ligase, and point mutants were generated by quick-change site-directed mutagenesis, using Phusion polymerase. All plasmids were verified by Sanger sequencing. The bacterial expression constructs for Mical1, MO-CH-LIM, MO-CH, and MO were previously described^52^. A detailed overview of all expression vectors employed in this study is presented in Supplementary Table 2.

### Recombinant protein expression and purification

Human Mical1 full-length and truncated C-terminal His6-tagged versions of MO, MO-CH, and MO-CH-LIM domains were expressed and purified with a modified protocol as described previously^52^. Proteins were recombinantly expressed in *E. coli* Rosetta (DE3) (Merck Millipore) cells in LB media supplemented with Ampicillin (125 µg/mL) and Chloramphenicol (34 µg/mL) and cells were grown at 37°C to OD_600_ nm = 0.8–1.0 and stored at 4°C for 30 min. Expression was induced by the addition of 0.5 mM IPTG, and cells were allowed to grow at 19°C for 14-16 hr. Cells were pelleted and stored at - 80°C until ready for purification. Cells were mechanically lysed by passing through a fluidizer (Microfluidic) in a buffer [Buffer A: 50 mM Hepes pH 8.0, 500 mM NaCl, 2 mM βME (2- Mercaptoethanol), 1 mM PMSF (Phenylmethylsulfonyl fluoride) and 1 mM CHAPS (3-((3- cholamidopropyl) dimethylammonio)-1-propanesulfonate)] and lysates were cleared by centrifugation at 75,600 g at 4 °C for 30 min using a JA 25.50 rotor (Beckman-Coulter). Next, the proteins were purified by Ni^2+^-affinity chromatography, and protein fractions were eluted with an imidazole gradient (Buffer B: buffer A supplemented with 500 mM imidazole pH 8.0). The His6-tagged protein was overnight dialyzed at 4 °C in a buffer (50 mM Tris pH 8.0, 100 mM NaCl, and 2 mM βME). Further purity was achieved by gel filtration (HiLoad Superdex 200 26/60, GE Healthcare) (Buffer: 20 mM Tris 8.0, 100 mM NaCl, and 2 mM DTE). Depending on purity, the protein was further purified by one to three successive rounds of anion exchange chromatography (POROS™ 50 HQ strong anion exchange resin; Thermo Scientific^TM^). The purified protein was collected, concentrated, and washed with final buffer (Storage buffer: 20 mM Tris pH 8.0, 100 mM NaCl, and 2 mM DTE) using Amicon^®^ Ultra-15 50K NMWL centrifugal filter devices; flash-frozen in liquid N_2_ and stored at −80 °C. Protein concentration was determined by using Bradford assay (Bio-Rad). Tag-free Mical1 full-length bMERB domain, truncated bMERB domains, and mutants were purified as described previously^28^.

Human Rab GTPases (tag-free) were expressed and purified as described previously^28^. Rabs were preparatively loaded with GppNHp (Guanosine-5’-[β-γ-Imido]-triphosphate) and the reaction was performed as described previously ^28^. Nucleotide exchange efficiency was quantified by C18 reversed-phase column (Prontosil C18, Bischoff Chromatography) with HPLC in 50 mM potassium phosphate buffer pH 6.6, 10 mM tetrabutylammonium bromide and 12% acetonitrile (v/v). Protein samples were heat precipitated at 95 °C for 5 min and centrifuged at 15700 g for 10 min and loaded (25 µM, 20 µL) on the column. Peaks were integrated and to determine the nucleotide retention times; a nucleotide standard run was performed. The purified protein was collected and concentrated; flash-frozen in liquid N_2_ and stored at −80 °C.

### Size exclusion chromatography coupled with multi-angle light scattering (SEC-MALS)

Full-length Mical1, MO-CH-LIM, MO-CH, and MO domains were analyzed using a Superdex 200 10/30 GL column coupled with a multi-angle light scattering (MALS) instrument (Wyatt Technology). The analysis was conducted in a buffer containing 20 mM Hepes pH 8.0, 50 mM NaCl, 1 mM MgCl_2_, and 1 mM TCEP at 25 °C. The bMERB domain was analyzed using a Superdex 75 10/30 GL column coupled with a MALS instrument (Wyatt Technology) in a buffer containing 20 mM Hepes pH 7.5, 50 mM NaCl, 1 mM MgCl_2_, and 1 mM TCEP. The chromatography system was connected in series with a light-scattering detector (Wyatt Dawn HELIOS II) and a refractive index detector (Wyatt Optilab t-rEX). BSA (Sigma) was used as a standard to calibrate the system, and 20 µL of each sample (2 mg/mL) was injected. Data analysis was performed with ASTRA 7.3.2 software (Wyatt Technology), yielding the molar mass and mass distribution (polydispersity) of the samples.

### Analytical size exclusion chromatography (aSEC)

The bMERB:GppNHp Rab (Rab8a_1-176_, Rab10_1-175_, Rab13_1-176_, and Rab15_1-176_) complex formation was analyzed by analytical size exclusion chromatography (aSEC). 110 µM of bMERB domain and 121 µM of GppNHp protein (Effector: Rab stoichiometry of 1: 1.1) were mixed in a buffer containing 20 mM Hepes pH 7.5, 50 mM NaCl, 1 mM MgCl_2,_ and 2 mM DTE and centrifuged for 15 min at 15700 g at 4°C. 40 µL of the mixture was injected into a Superdex 75 10/300 GL gel filtration column (GE Healthcare) pre-equilibrated with the Rab buffer with a flow rate of 0.5 mL/min at room temperature and absorption at 280 nm was recorded.

### Isothermal titration calorimetry (ITC)

Protein-protein interaction measurements were conducted by isothermal titration calorimetry (ITC) using an ITC200 and PEAQ-ITC microcalorimeter (MicroCal). Mical1_fl_/bMERB:Rab interaction measurements were performed in the buffer containing 20 mM Hepes 7.5, 50 mM NaCl, 1 mM MgCl_2,_ and 1 mM Tris (2-carboxymethyl) phosphine (TCEP) whereas MO-CH-LIM/MO-CH/MO:bMERB interactions were performed in buffer containing 20 mM Hepes 8.0, 150 mM NaCl and 1 mM TCEP at 25°C. Wild types and mutant proteins were dialyzed overnight in their respective buffer. Samples were centrifuged at 15700 g for 30 min at 4°C and protein concentration was determined by Bradford assay (Bio-Rad). 600 µM of GppNHp Rab8a_1-176_ was titrated into the cell containing 30 µM bMERB domain and for full-length Mical1: Rab8 measurement, 800 µM of GppNHp Rab8a_1-176_ was titrated into the cell containing 40 µM full length Mical1. For MO-CH-LIM/MO-CH/MO:bMERB interaction 600 µM of bMERB domain was titrated into the cell containing 30 µM MO-CH-LIM/MO-CH/MO domain. For the control experiments, the buffer was titrated into the cell containing the protein of interest. In the second control experiment, the protein in the syringe was titrated against the buffer. The binding isotherms were integrated and corrected for the offset by subtracting the control experiment, and the data were fitted to a one-site-binding or two-site-binding model using PEAQ-ITC software. The reported ITC result is representative of one of at least three independent measurements.

### AlphaFold2 molecular modeling

The Mical1 models (Mical1_fl_ and MO-CH-LIM:bMERB complex) were generated using AlphaFold multimer v2.3.1^31^.

### Cross-linking experiment

Full-length Mical1 (25 µM) was mixed with a 10-fold excess of freshly dissolved DSBU (disuccinimidyl dibutyric urea; Thermo Fisher SCIENTIFIC, 50 mM stock) in DMSO in a cross-linking buffer (20 mM Hepes, 100 mM NaCl, and 2 mM DTE). In a second experiment, 50 µM of MO-CH-LIM (residues 1-781) was mixed with 100 µM of bMERB domain (residues 918-1067) in a final ratio of 1:2, respectively, and again, 10 times DSBU was added in the crosslinking buffer. Samples were incubated at room temperature for 1.30 hours, and the crosslinking reactions were quenched by the addition of 100 mM Tris, pH 8.0. The samples were flash-frozen for further processing, as described previously^53^. Briefly, the processing steps included the reduction of disulfide bonds using tris(2-carboxyethyl) phosphine, alkylation of free thiol groups on cysteines with iodoacetamide, and proteolysis with trypsin. The resulting peptides were enriched as described previously^54^. Briefly, the peptides were fractionated using a Superdex™ 30 Increase 3.2/300 column in a buffer containing 50 mM ammonium acetate, 30% acetonitrile, and 0.1% formic acid. Eluted samples were dried using a centrifugal evaporator and stored at −20 °C until measurement.

### LC-MS/MS analysis

LC-MS/MS analysis was performed as described previously^54^. Briefly, peptides were dissolved in 20 μL of water containing 0.1% TFA. A total of 3 μL of this peptide solution were separated on an Ultimate 3000 RSLC nano system (precolumn: C18, Acclaim PepMap, 300 μm × 5 mm, 5 μm, 100 Å, separation column: C18, Acclaim PepMap, 75 μm × 500 mm, 2 μm, 100 Å, Thermo Fisher Scientific). After loading the sample on the pre-column, a multistep gradient from 5−40% B (90 min), 40−60% B (5 min), and 60−95% B (5 min) was used with a flow rate of 300 nL/min; solvent A: water + 0.1% formic acid; solvent B: acetonitrile + 0.1% formic acid. The nano-HPLC system was coupled to a Q-Exactive Plus mass spectrometer (Thermo Fisher Scientific). Data were acquired in data-dependent MS/MS mode. For full scan MS, we used a mass range of m/z 300−1800, resolution of R = 140000 at m/z 200, one microscan using an automated gain control (AGC) target of 3e6, and a maximum injection time (IT) of 50 ms. Then, we acquired up to 10 HCD MS/MS scans of the most intense at least doubly charged ions (resolution 17500, AGC target 1e5, IT 100 ms, isolation window 4.0 m/z, normalized collision energy 25.0, intensity threshold 2e4, dynamic exclusion 20.0 s). All spectra were recorded in profile mode.

### Data analysis and cross-link identification

Raw data from the Q-Exactive were converted to MGF (Mascot generic files) format with the program msConvert GUI from ProteoWizard Toolkit version 3^55^ and the mgf-files of all fractions of a crosslink were merged into one file. Program MeroX version 2.0.1.4^56,57^ was used for cross-link identification. MS data in MGF format and protein sequences in FASTA format were loaded on the program and the cross-linked residues were searched. In the settings, the precursor precision and the fragment ion precision were set to 10.0 and 20.0 ppm, respectively. MS1 data were re-calibrated by 5 ppm according to the average mass deviation of the raw MS data. The FDR was set to 50, i.e. no filtering according to FDR with MeroX was done. RISEUP mode was used, and the maximum missing ions was set to 1. Crosslinks data were exported to XiViewer^58^.

### Crystallization and structure determination

Initial crystallization condition screens for the protein complexes described in the paper were performed using the JSG CORE I-IV, Pact, and Protein Complex suites (Qiagen). The sitting-drop vapor diffusion method was employed, with a reservoir volume of 70 μL and a drop volume of 0.1 μL for the protein (300 µM complexes, 1:1 Rab:effector) and 0.1 μL for the reservoir solution at 20°C. The best conditions were then optimized using the sitting-drop vapor diffusion method, with varying drop sizes, to obtain well-diffracting crystals. The complex of bMERB_V978A_V985A_:Rab10_1-175_ (300 µM of 1:1 complex) was crystallized in 0.17 M Sodium acetate, 0.085 M Tris-HCl pH 8.5, 25.5% (w/v) PEG 4000 and 15% (v/v) glycerol. The complex bMERB_V978A_:Rab10_1-175_ (300 µM of 1:1 complex) was crystallized in 0.1 M Imidazole pH 8.0, 5% (w/v) PEG 3000 and 30% (v/v) PEG 200. Crystals were fished directly from the crystallization drop and flash-cooled in liquid nitrogen. Diffraction data of bMERB_V978A_V985A_:Rab10_1-175_ was collected at 100 K on beamline X10SA at the Swiss Light Source (Paul Scherrer Institute, Villigen, Switzerland). For the bMERB_V978A_V985A_:Rab10_1-175_ complex crystal, a native data set was collected at a wavelength of 0.999968 Å. Whereas diffraction data of bMERB_V978A_:Rab10_1-175_ was collected at 100 K on beamline ID23-2 at the European Synchrotron Radiation Facility (Grenoble, France). For the bMERB_V978A_:Rab10_1-175_ complex crystal, a native data set was collected at a wavelength of 0.873130 Å. Data were integrated and scaled with XDS^59^.

The crystal of the bMERB_V978A_V985A_:Rab10_1-175_ complex diffracted to a resolution of 1.8 Å (space group P2_1_ with a = 53.673 Å, b = 48.807 Å, c = 79.199 Å), and a single copy of the complex is present in the asymmetric unit of the crystal. The initial model for the bMERB_V978A_V985A_:Rab10_1-175_ complex was obtained by molecular replacement using PHASER^60^ with the crystal structure of the Mical1_bMERB_:Rab 10 (PDB: 5LPN) as a search model^28^. The partial model was completed by manual building in Coot^61^. For the bMERB_V978A_:Rab10_1-175_ complex, the crystal diffracted to a resolution of 2.05 Å (space group P2_1_ with a = 53.517 Å, b = 49.434 Å, c = 79.095 Å), and one copy of the complex constitutes the asymmetric unit of the crystal. The partial model was obtained by molecular replacement using PHASER^60^ and the bMERB_V978A_V985A_:Rab10_1-175_ was used as a search model. The initial model was completed by manual building in Coot^61^. The final models were refined to convergence with phenix.refine^62^.

Data collection and refinement statistics are summarized in Supplementary Table 1. Structural figures were prepared using PyMOL (DeLano Scientific; http://www.pymol.org).

### Actin co-sedimentation assay

Rabbit skeletal muscle G-actin (AKL99) was purchased from Cytoskeleton. Inc and polymerized into F-actin according to the manufacturer’s protocol. F-actin (10 μM) was incubated for 1 hr at room temperature (RT) with MO/MO-CH/MO-CH-LIM/Mical1_fl_ (10 µM) in a buffer containing 20 mM Hepes (pH 8.0), 50 mM KCl, 2 mM MgCl_2_, 1 mM DTT, and 2 mM NaN_3_. Samples were centrifuged at 100,000 g for 1 hr at 4°C. The supernatant and pellet were subjected to 4-15% gradient SDS-PAGE, followed by Coomassie Brilliant Blue staining.

### Bioinformatics

Multiple sequence alignments were generated using Clustal Omega^63^. The protein interaction interfaces from the asymmetric unit were examined in detail using the PDBePISA server (Proteins, Interfaces, Structures, and Assemblies)^64^. The DALI server was used for structural comparison^65^.

### Data availability

The plasmids generated in this study are available from the corresponding author upon request. Protein coordinates and structure factors have been submitted to the Protein Data Bank under accession codes.

PDB: 9G0C (bMERB_V978A_V985A_:Rab10)

PDB: 9G0D (bMERB_V978A_:Rab10)

Source data are provided with this paper.

## Acknowledgments

We are grateful to Prof. Maria Vanoni for sharing the full-length Mical1, MO-CH-LIM, MO-CH, and MO bacterial expression constructs and providing the purification protocol. We also thank the beamline staff of the Swiss Light Source (SLS) X10SA at the Paul Scherrer Institute, Villigen, Switzerland, and the beamline staff of the European Synchrotron Radiation Facility at Grenoble, France, for their assistance during data collection. We thank Dr. Raphael Gasper and Dr. Matthias Müller for the X-ray diffraction data collection. We express our gratitude to Petra Geue for conducting the SEC-MALS measurements and data analysis. Special thanks are extended to Franziska Müller for her contributions to the size exclusion chromatography experiment for the peptides. We are thankful to Malte Metz and Andreas Brockmeyer for conducting the LC-MS/MS runs and method setup. Additionally, we acknowledge financial support from the Max-Planck-Society and the Deutsche Forschungsgemeinschaft (grant GO 284/10-1 to R.S.G). Open access funding provided by Max Planck Society.

## Author contributions

A.R. conceived and designed the study, carried out protein expression and purification, conducted all biochemical studies, cross-linking experiments, crystallization, and determined X-ray structures. P.J. performed LC-MS/MS experiments, conducted data analysis, and identified cross-links. I.R.V. conducted Mical1 AlphaFold2 modeling and analyzed the structure. R.S.G. acquired funding. A.R., P.J., I.R.V., and R.S.G. analyzed and interpreted the data. A.R. wrote the manuscript with critical input from all authors.

