## Supplementary Information for "Biochemical and structural insights into the auto-inhibited state of Mical1 and its activation by Rab8"

Rai *et al*.,

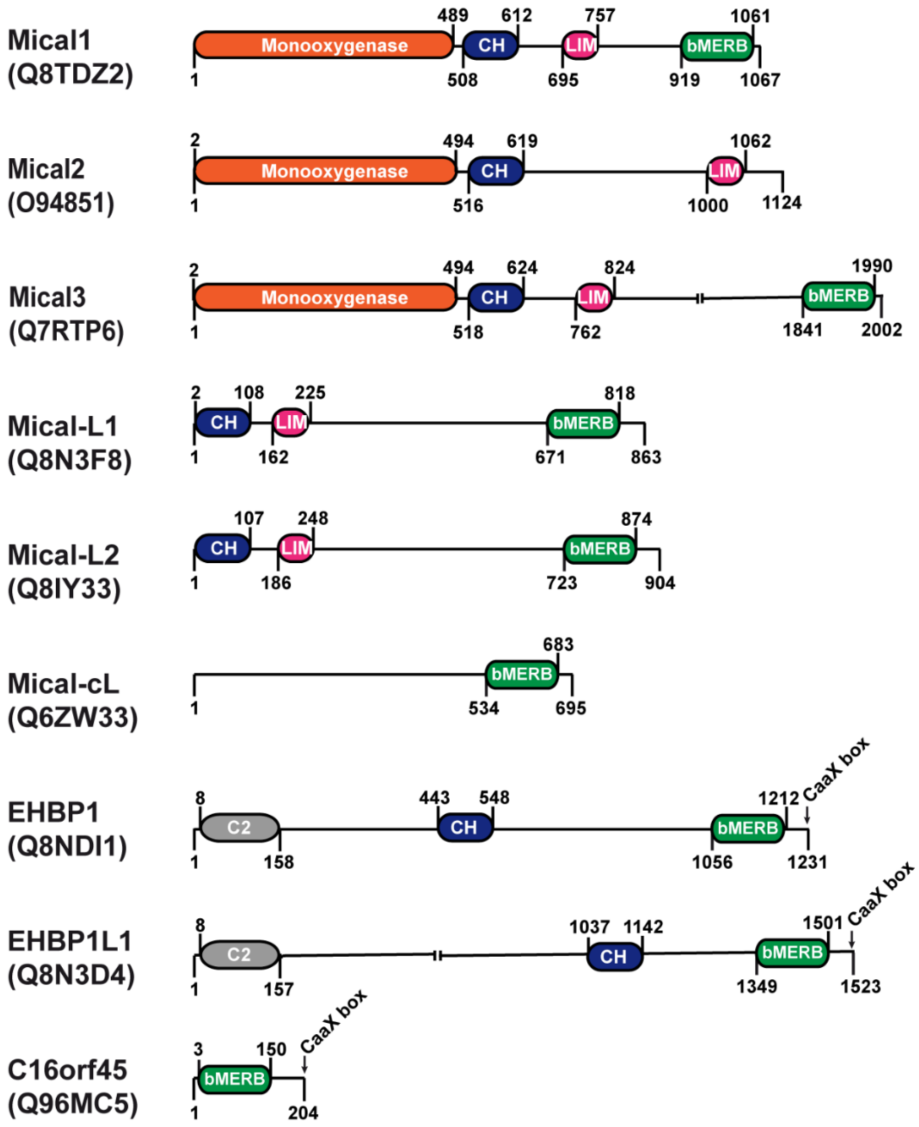


**Supplementary Fig. 1: Domain organization of bMERB domain-containing proteins.** Micals contain an N-terminal Monooxygenase (orange), a CH (blue), a LIM domain (magenta), and a C-terminal bMERB (green) domain, whereas Mical-like proteins lack the MO domain. Along with the CH and bMERB domains, EHBPs also contain an N-terminal lipid-interacting C2 domain and a C-terminal CaaX box (prenylation motif). Mical-cL and C16orf45 have only the bMERB domain. Domain boundaries are taken from UniProt IDs.

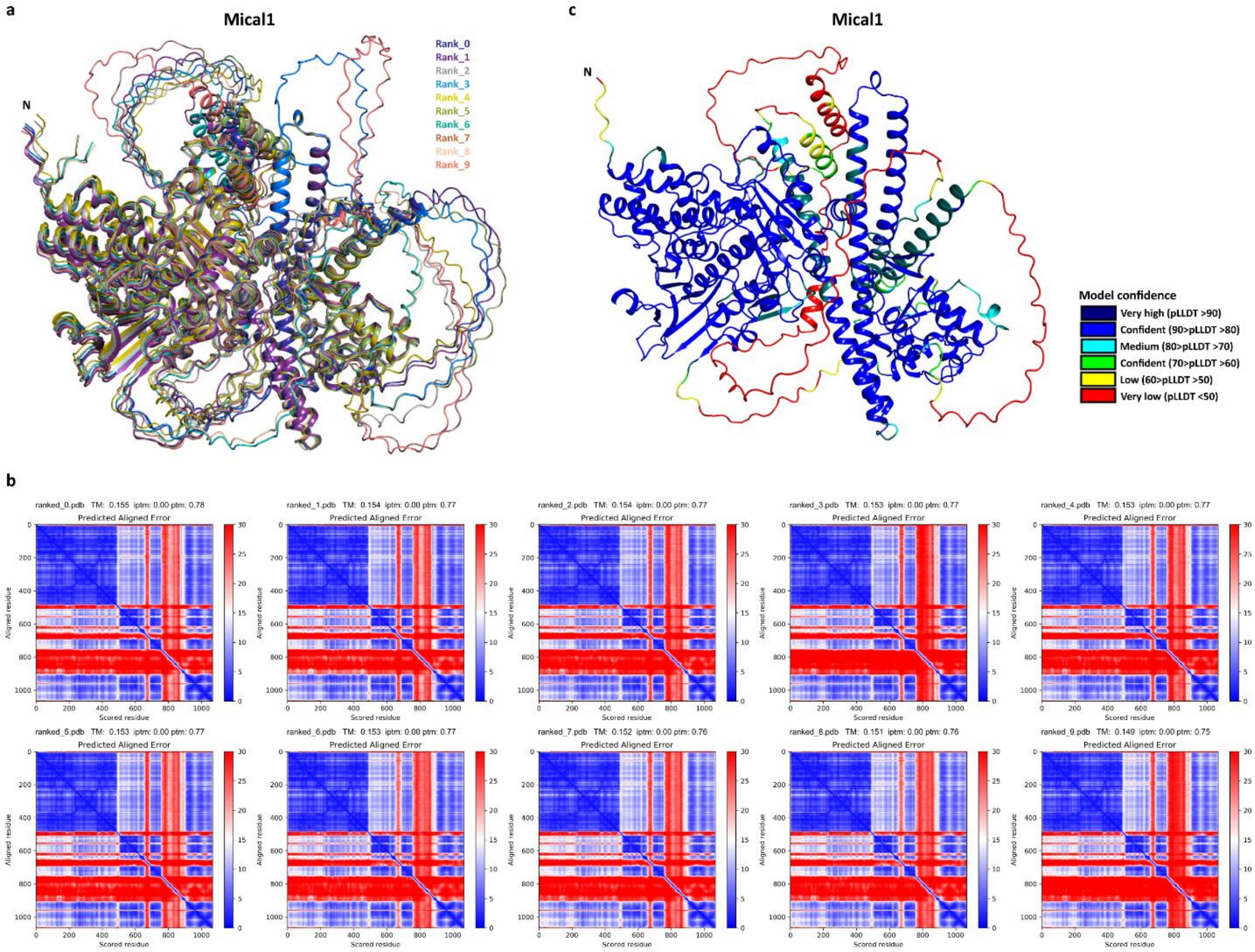


**Supplementary Fig. 2: Mical1 auto-inhibited AlphaFold2 model confidence. (a)** Overlay of AlphaFold2[1] predicted Mical1 models. **(b)** Predicted aligned error (PAE) scores for distance error of the top-ranked. **(c)** Cartoon presentation of AlphaFold2 predicted top model is color-coded based on predicted local difference distance test (pLDDT) scores.

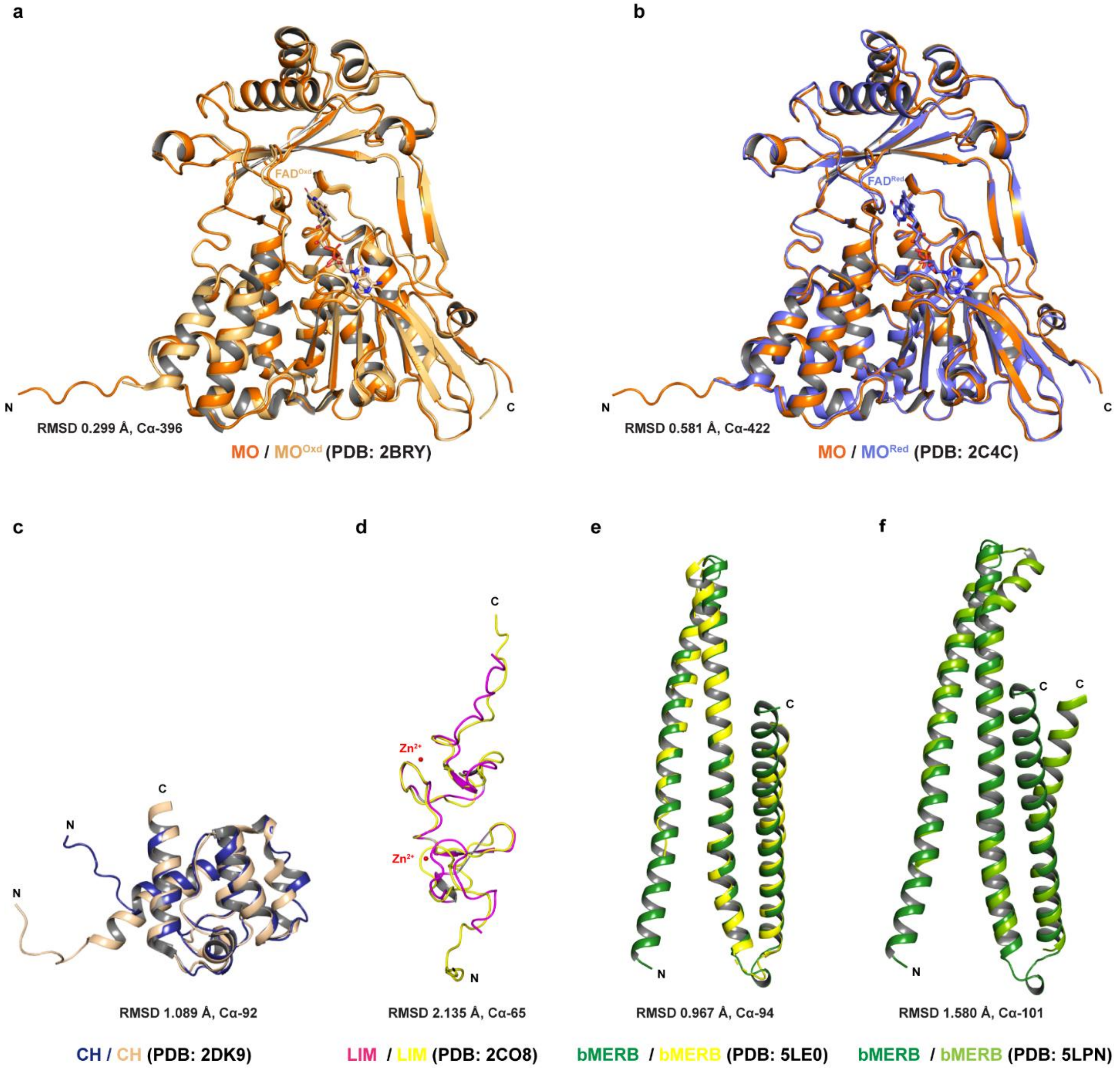


**Supplementary Fig. 3: Structural superposition of the Mical1 domains in the auto-inhibited state with other published Mical1 domain structures. (a-b)** Structural overlay of the MO domain with the oxidized mouse Mical1 MO domain (light orange, PDB: 2BRY)[2] and the reduced mouse Mical1 MO domain (slate, PDB: 2C4C)[2]. **(c)** Structural overlay of the CH domain with the NMR structure of the Mical1 CH domain[3]. **(d)** Structural superposition of the NMR domain with the NMR structure of the Mical1 LIM domain[4]. **(e-f)** Structural superposition of the bMERB domain with the isolated bMERB domain[5] and with the bMERb domain of Rab10 structure[6].

```
hMical1    918    KEEEMKRFCKAQTIQRRLNEIEAALRELEAEGVKLELALRRQSSSPEQQKKLWVGQLLQL    977
rMical1    905    KEEEMKRFCKAQAIQRRLNEIEAAMRELETEGMKLEVALRKESSSPEKQKKLWLEQLLQL    964
mMical1    905    KEEEMKRFCKAQAIQRRLNEIEATMRELEAEGTKLELALRKESSSPEQQKKLWLDQLLRL    964
bMical1    921    KEEEMKRFCKAQAIQRRLNEIEAALRELEARGTELELALRSQSSSPEKQKALWVEQLLQL    980
gMical1    937    KEEEMKRFCKAQTIQRRLNEIEAALRELEAEGVKLELALRRQSSSPEQQKKLWVGQLLQL    996
cMical1    846    REEEMKRFCKAQAVQRRLNEIETALRELEAEGVKLELALRSQSTSPEQQKTLWLEQLLQL    905
                  :***********::********:::****:.* :**:*** :*:***:** **: ***:*

hMical1    978    VDKKNSLVAEEAELMITVQELNLEEKQWQLDQELRGYMNREENLKTAADRQAEDQVLRKL    1037
rMical1    965    IQKKNSLVTEEAELMITVQELDLEEKQRQLDHEFRGI-NREETLKTQADRLSEDRVLRKL    1023
mMical1    965    IQKKNSLVTEEAELMITVQELDLEEKQRQLDHELRGYMNREETMKTEADLQSENQVLRKL    1024
bMical1    981    VQKKNSLVAEEAELMITVQELNLEEKQWQLDQELRTYMNREETLKTAADRQAEDQVLRKL    1040
gMical1    997    VDKKNSLVAEEAELMITVQELNLEEKQWQLDQELRGYMNREETLKTAADRQAEDQVLRKL    1056
cMical1    906    VEKKNSLVAEEAELMITVQELKLEDQQWQLDQELRGYMNREEFLKTPADRQAEDQVLRKL    965
                  ::******:************.**::* ***:*:*   **** :** **  :*::*****

hMical1    1038 VDLVNQRDALIRFQEERRLSELALGTGAQG-    1067
rMical1    1024 LDVVNQRDALIQFQEERRLREMPV-------    1047
mMical1    1025 LEVVNQRDALIQFQEERRLREMPA-------    1048
bMical1    1041 LDVVNQRDALIRLQEERRLSELASEPGVQG-    1070
gMical1    1057 VDLVNQRDALIRFQEERRLSELALGTGAQG-    1086
cMical1    966  LDVVNQRDALIRFQEECRLSELASEPGAQG-    995
                :::********::*** ** *:
```

**Supplementary Fig. 4:** Sequence alignment of human Mical1 bMERB domain with mammalian homologs from *Mus musculus* (mMical1), *Rattus norvegicus* (rMical1), *Bos taurus* (bMical1), *Gorilla gorilla gorilla* (gMical1) and *Canis lupus familiaris* (cMical1) using Clustal Omega[7]. Identical residues are denoted by *, strongly similar residues by: and . indicates weakly similar residues. Residues involved in Mical MO-CH-LIM domain interactions are labeled in red.

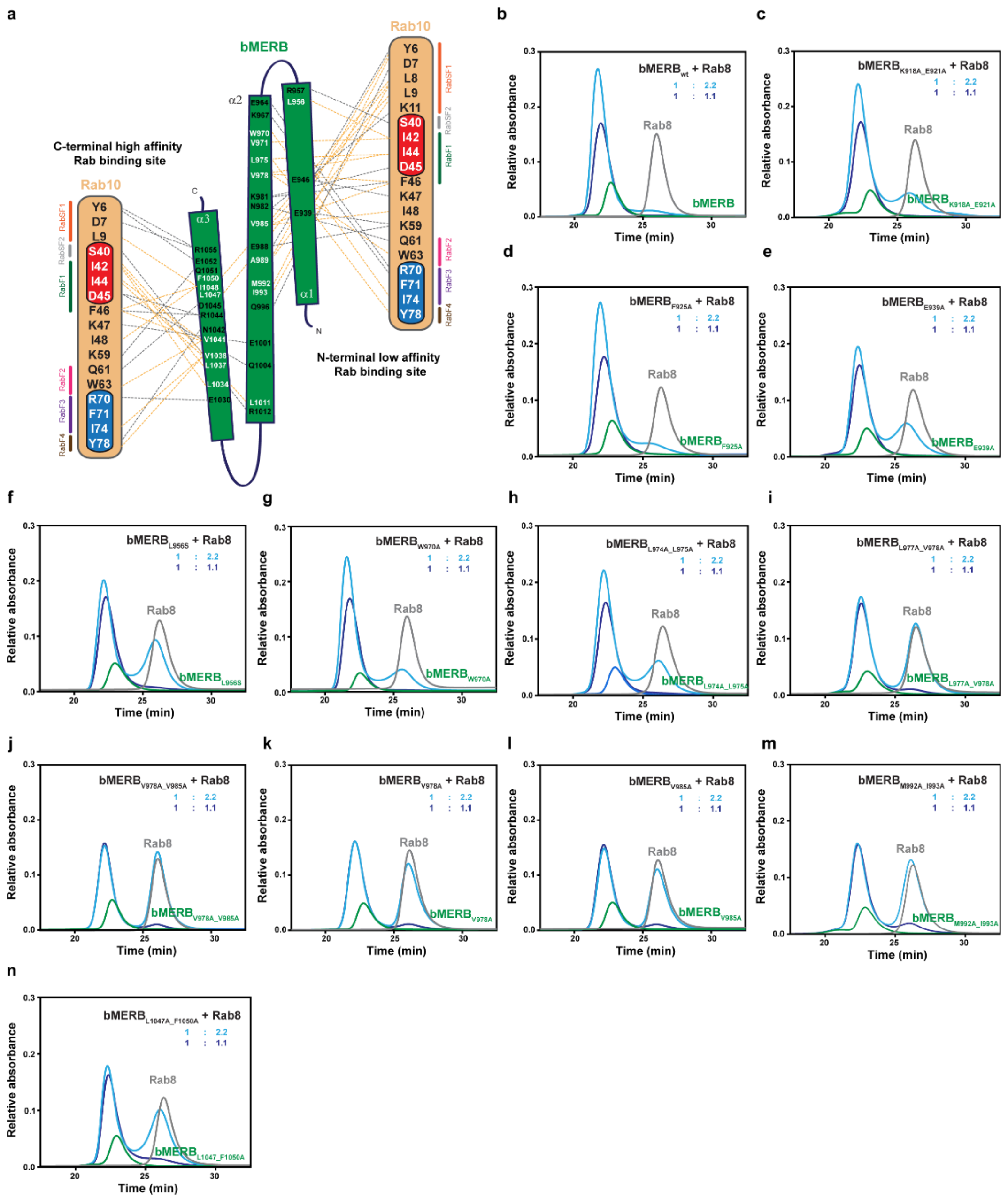

**Supplementary Fig 5. Interaction of Mical1 bMERB RBS1 mutants with Rab8. (a)** Schematic illustration of the interactions between the bMERB domain and Rab10 (PDB: 5LPN)[6]. Hydrogen bonds and ionic interactions are shown in gray dashed lines, and light orange dashed lines indicate hydrophobic interactions. RabSF1, RabSF2, RabF1, RabF2, RabF3, and RabF4 motifs are shown in orange, gray, green, pink, purple, and brown respectively. **(b-n)** The binding of Rab8 (gray, 121/242 µM) with different RBS1 bMERB mutants (green, 110 µM) was systematically tested on a Superdex75 10/300 GL column. Mutants K918A_E921A, F925A, and W970A form clear complexes, and an increase in the concentration of Rab8 leads to a more complex formation, suggesting a 1:2 bMERB: Rab stoichiometry. Conversely, in the case of E939A, L956S, and L974A_L975A, a small increase in complex formation is observed upon increasing the Rab8 concentration, suggesting the mutation leads to a decrease in binding affinity at RBS1. No change in complex formation upon increasing Rab8 concentration was observed for the mutants L977A_V978A, V978A_V985A, V978A, V985A, M992A_I993A, and L1047A_F1050A, suggesting single Rab binding (1:1 stoichiometry) or lower affinity for both Rabs. The data are representative of at least three repetitions.

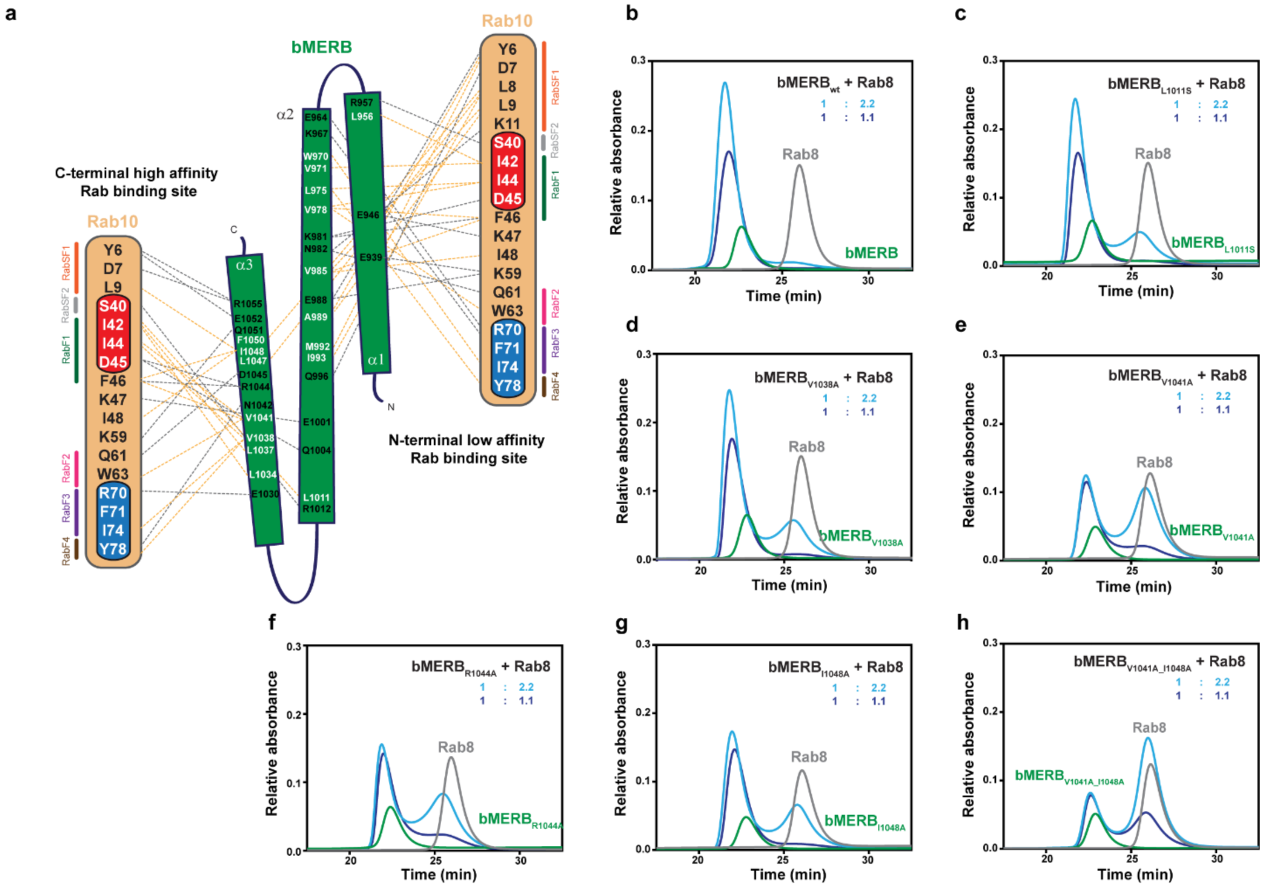


**Supplementary Fig 6. Interaction of Mical1 bMERB RBS2 mutants with Rab8. (a)** Schematic illustration of the interactions between the bMERB domain and Rab10 (PDB: 5LPN)[6]. Hydrogen bonds and ionic interactions are shown in gray dashed lines, while light orange dashed lines indicate hydrophobic interactions. RabSF1, RabSF2, RabF1, RabF2, RabF3, and RabF4 motifs are shown in orange, gray, green, pink, purple, and brown respectively. **(b-h)** The binding of Rab8 (gray, 121/242 µM) with different RBS2 bMERB mutants (green, 110 µM) was systematically tested on a Superdex75 10/300 GL column. Mutants L1011S and V1038 form clear complexes, and an increase in the concentration of Rab8 leads to a more complex formation, suggesting a 1:2 bMERB:Rab stoichiometry. However, compared to the wild-type protein, no change in complex formation upon increasing Rab8 concentration was observed for the mutants V1041A, R1044A, and I1048A, suggesting a single Rab binding (1:1 stoichiometry) or lower Rab binding affinity for both sites. The bMERB construct V1041A_I1048A failed to form stable complexes with Rab8, suggesting that these mutations at the C-terminal RBS2 also affect the binding at the N-terminal. The data are representative of at least three repetitions.

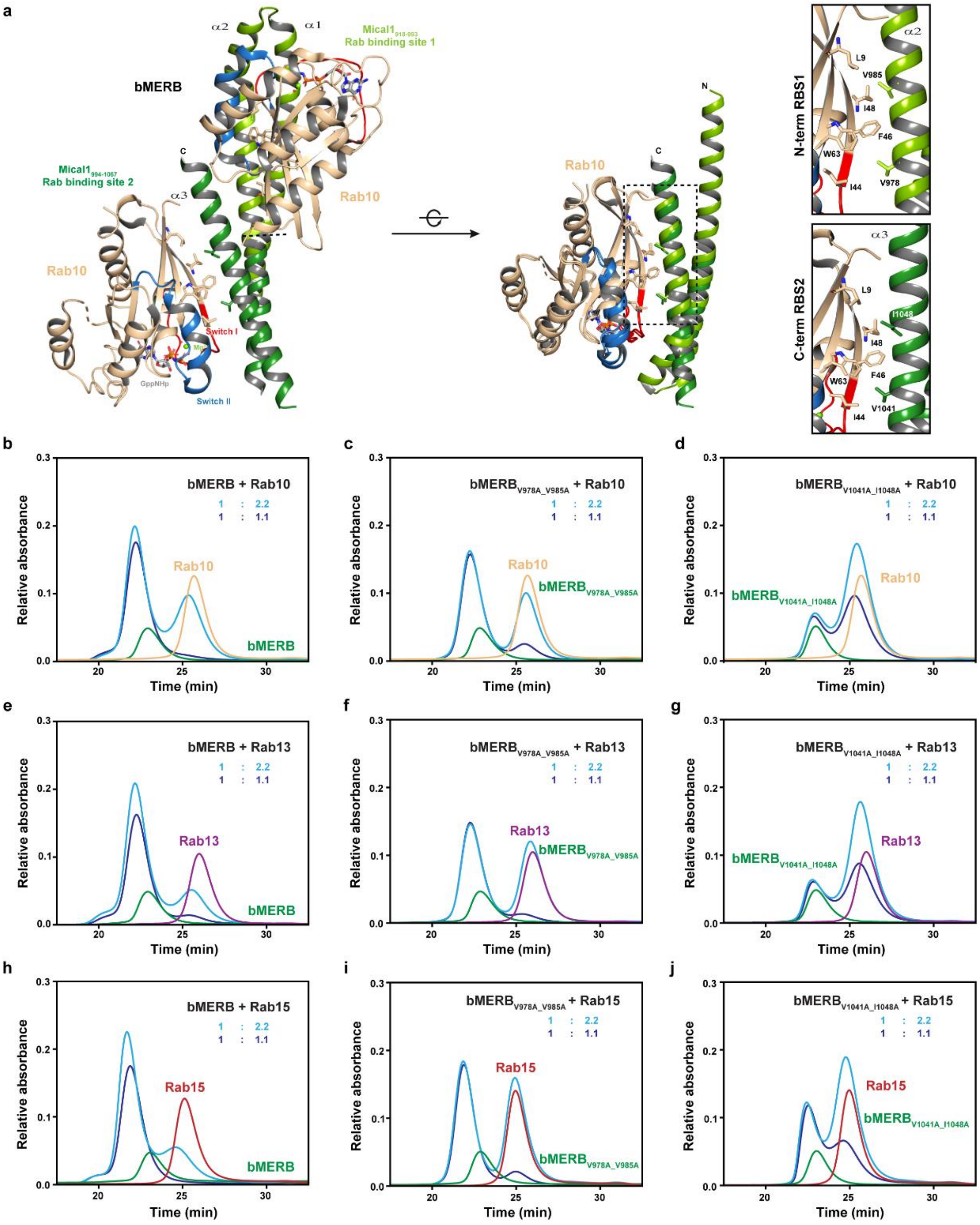


**Supplementary Fig 7. Interaction of Mical1 bMERB mutants with Rab10/Rab13/Rab15**. **(a)** Cartoon representation of Mical1 bMERB:Rab10 complex structure (PDB: 5LPN)[6]. Structural alignment of the N- and C-terminal halves of the Mical1 bMERB domain shows strong conservation between both Rab8 binding sites. V978/V1041 form hydrophobic interactions with I44, F46, and W63 of Rab10, whereas V985/I1048 form hydrophobic interactions with L9, F46, and I48 of Rab10. **(b-j)** The binding of Rab10 (wheat, 121/242 µM), Rab13 (purple, 121/242 µM), and Rab15 (red, 121/242 µM) with RBS1/RBS2 bMERB mutants (green, 110 µM) was systematically tested on a Superdex75 10/300 GL column. Compared to the wild-type protein, mutant V978A_V985A shows no increase in complex formation upon an increase in Rab concentration, suggesting that the mutation at the N-terminal leads to the loss of Rab binding at the N-terminal binding site. In contrast, mutant V1041A_I1048A shows no stable complex formation with Rab10/Rab13/Rab15. The data are representative of at least three repetitions.

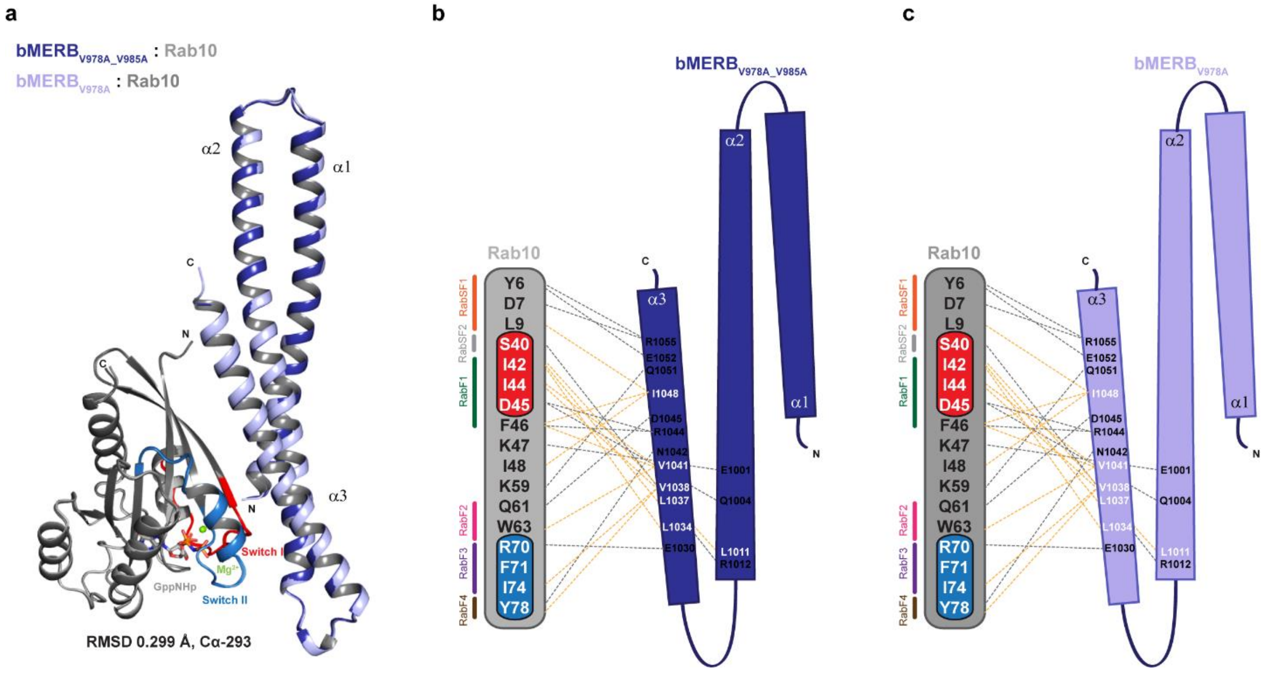


**Supplementary Fig 8. Structure of Mical1 bMERB$_{V978A_V985A}$:Rab8 and Mical1 bMERB$_{V978A}$:Rab8 complex. (a)** An overlay of the bMERB$_{V978A_V985A}$:Rab8 complex structure with the bMERB$_{V978A}$:Rab8 complex. **(b-c)** Schematic illustration of the interactions between the bMERB$_{V978A_V985A}$/bMERB$_{V978A}$ domain and Rab8. Hydrogen bonds and ionic interactions are shown in gray dashed lines, while light orange dashed lines indicate hydrophobic interactions. RabSF1, RabSF2, RabF1, RabF2, RabF3, and RabF4 motifs are shown in orange, gray, green, pink, purple, and brown, respectively.

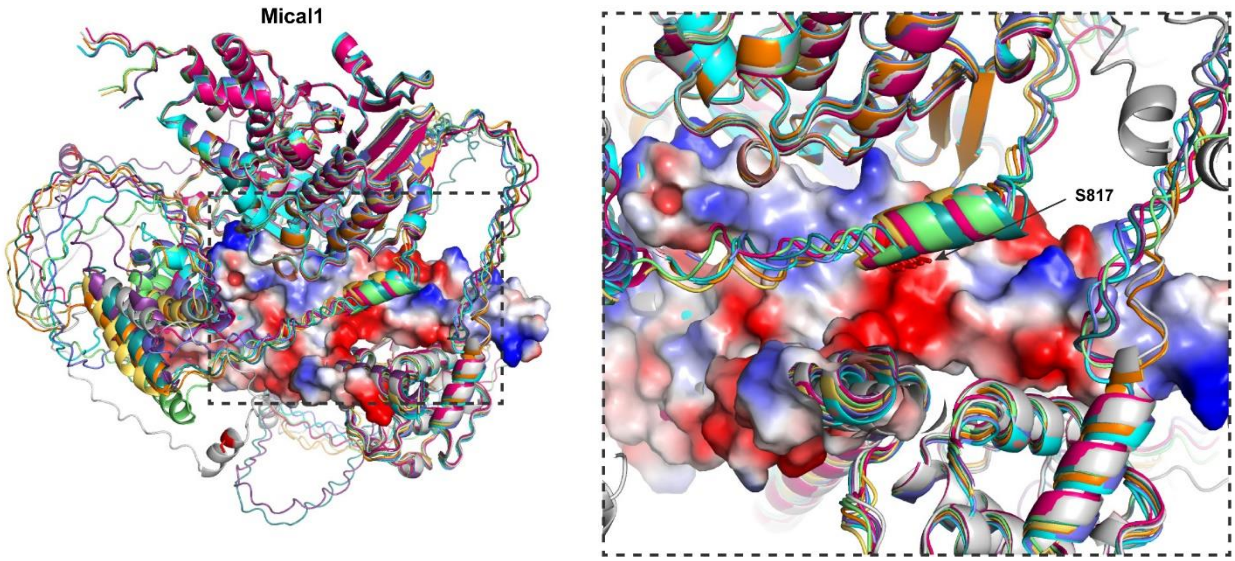


**Supplementary Fig 9. Schematic diagram of Mical1 AlphaFold2 model.** Overlay of AlphaFold2 predicted Mical1 models with the surface electrostatic potential of the bMERB domain. The inset shows the PAK1 phosphorylation site S187, with the side chain of S187 pointing towards a negatively charged patch of the Mical1 bMERB domain.

**Supplementary Table 1: Data-collection and refinement statistics (values in parentheses are for the outer shell).**

| | $bMERB_{V978A\_V985A}$:Rab10 | $bMERB_{V978A}$:Rab10 |
|---|---|---|
| **Data collection#** | | |
| X-Ray Source | X10SA SLS | ESRF-ID23-3 |
| Wavelength (Å) | 0.999968 | 0.873130 |
| Resolution range (Å) | 41.22-1.80 (1.85-1.80) | 36.29-2.05 (2.10-2.05) |
| Space group | $P2_1$ | $P2_1$ |
| Unit cell<br>a, b, c (Å)<br>α, β, γ (°) | <br>53.39, 48.56, 78.77<br>90.0, 97.981, 90.0 | 53.77, 49.65, 79.33<br>90.0, 98.538, 90.0 |
| Total reflections | 258299 (17871) | 176980 (13279) |
| Unique reflections | 37165 (2758) | 26092 (1923) |
| Multiplicity | 6.9 (6.47) | 6.78 (6.9) |
| Completeness (%) | 99.5 (99.7) | 99.3 (99.7) |
| Mean I/sigma(I) | 17.93 (1.85) | 11.36 (1.8) |
| Rmerge (%) | 5.1 (72.7) | 9.5 (133.8) |
| $R_{meas}$ (%) | 5.5 (79.1) | 10.4 (144.2) |
| CC1/2 | 99.9 (92.5) | 99.7 (79.9) |
| **Refinement** | | |
| Resolution range (Å) | 41.22-1.80 (1.85-1.80) | 36.29-2.05 (2.10-2.05) |
| Reflections used in refinement | 37089 | 26058 |
| Reflections used for R-free | 1854 | 1304 |
| $R_{work}$ | 0.1928 (0.3356) | 0.2032 (0.2973) |
| $R_{free}$ | 0.2312 (0.3435) | 0.2487 (0.2957) |
| Total number of atoms | 2829 | 2641 |
| R.m.s deviations<br>Bond length (Å)<br>Bond angles (°) | <br>0.006<br>0.83 | <br>0.007<br>0.808 |
| Ramachandran plot<br>Favored (%)<br>Additionally allowed (%)<br>Outliers (%) | <br>98.07<br>1.93<br>0.0 | <br>96.79<br>2.56<br>0.64 |
| B-factors ($Å^2$)<br>Protein<br>Ligands<br>Water | <br>47.68<br>42.54<br>54.70 | <br>58.05<br>52.59<br>54.69 |
| PDB ID | 9G0C | 9G0D |

**Supplementary Table 2: Expression constructs used in this study.**

| Construct | Plasmid | Description | Insert boundaries (restriction sites) | Purification | Purpose |
|---|---|---|---|---|---|
| Mical1fl_c-6His | pET23a-hs-Mical1fl Cter.His | Vitali *et al*., 2016[8] Hs-Mical1fl-6His | 1-1067 aa (NdeI-XhoI) | Ni-NTA, Superdex 200, IEX | SEC-MALS, ITC, Crystallization, actin co-sedimentation |
| MO-CH-LIM_c-6HIs | pET23a-hs-MO-CH-LIM Cter.His | Vitali *et al*., 2016[8] Hs-$Mical1_{1\text{-}781}$-6His | 1-781 aa (NdeI-XhoI) | Ni-NTA, Superdex 200, IEX | SEC-MALS, ITC, Crystallization, actin co-sedimentation |
| MO-CH_c-6HIs | pET23a-hs- MO-CH cter.His | Vitali *et al*., 2016[8] Hs-$Mical1_{1\text{-}615}$-6His | 1-615 aa (NdeI-XhoI) | Ni-NTA, Superdex 200, IEX | SEC-MALS, ITC, actin co-sedimentation |
| MO_c-6HIs | pET23a-hs-MO cter.His | Vitali *et al*., 2016[8] Hs-$Mical1_{1\text{-}488}$-6His | 1-488 aa (NdeI-XhoI) | Ni-NTA, Superdex 200, IEX | SEC-MALS, ITC, actin co-sedimentation |
| bMERB | pET19mod-hs-Mical1_918-1067 | Rai *et al*., 2016[6] 6His-TEV-hs-$Mical1_{918\text{-}1067}$ | 918-1067 aa (NdeI-XhoI) | Ni-NTA, TEV, Ni-NTA, Superdex 75 | SEC-MALS, ITC, SEC-MALS, ITC, Crystallization |
| $bMERB_{\_H1\text{-}2}$ | pET19mod-hs-Mical1 918-1020 | Rai *et al*., 2016[6] 6His-TEV-hs-$Mical1_{918\text{-}1020}$ | 918-1020 aa (NdeI-XhoI) | Ni-NTA, TEV, Ni-NTA, Superdex 75 | ITC |
| $bMERB_{\_H2\text{-}3}$ | pET19mod-hs- Mical1 960-1067 | Rai *et al*., 2016[6] 6His-TEV-hs-$Mical1_{960\text{-}1067}$ | 960-1067aa (NdeI-XhoI) | Ni-NTA, TEV, Ni-NTA, Superdex 75 | ITC |
| $bMERB_{K918A\_E921A}$ | pET19mod -hs-Mical1_$bMERB_{K918A_E921A}$ | 6His-TEV-hs-$Mical1_{918\text{-}1067\_K918A\_E921A}$ | 918-1067 aa (NdeI-XhoI) Quick change to mutate K918 and E921 to A | Ni-NTA, TEV, Ni-NTA, Superdex 75 | ITC |
| $bMERB_{F925A}$ | pET19mod -hs-Mical1_$bMERB_{F925A}$ | 6His-TEV-hs-$Mical1_{918\text{-}1067\_F925A}$ | 918-1067 aa (NdeI-XhoI) Quick change to mutate F925 to A | Ni-NTA, TEV, Ni-NTA, Superdex 75 | ITC |
| $bMERB_{E939A}$ | pET19mod-hs-Mical1_$bMERB_{E939A}$ | 6His-TEV-hs-$Mical1_{918\text{-}1067\_E939A}$ | 918-1067 aa (NdeI-XhoI) Quick change to mutate E939 to A | Ni-NTA, TEV, Ni-NTA, Superdex 75 | ITC |
| $bMERB_{L974A\_L975A}$ | pET19mod-hs-Mical1_$bMERB_{L974A_L975A}$ | 6His-TEV-hs-$Mical1_{918\text{-}1067\_L974A\_L975A}$ | 918-1067 aa (NdeI-XhoI) Quick change to mutate L974 and L975 to A | Ni-NTA, TEV, Ni-NTA, Superdex 75 | ITC |
| $bMERB_{L977A\_V978A}$ | pET19mod-hs-Mical1_$bMERB_{L977A_V978A}$ | 6His-TEV-hs-$Mical1_{918\text{-}1067\_L977A\_V978A}$ | 918-1067 aa (NdeI-XhoI) Quick change to mutate L977 and V978 to A | Ni-NTA, TEV, Ni-NTA, Superdex 75 | ITC |
| $bMERB_{M992A\_I993A}$ | pET19mod-hs-Mical1_$bMERB_{M992A_I993A}$ | 6His-TEV-hs-$Mical1_{918\text{-}1067\_M992A\_I993A}$ | 918-1067 aa (NdeI-XhoI) Quick change to mutate M992 and I993 to A | Ni-NTA, TEV, Ni-NTA, Superdex 75 | ITC |
| $bMERB_{L1047A\_L1050A}$ | pET19mod-hs-Mical1_$bMERB_{L1047A_L1050A}$ | 6His-TEV-hs-$Mical1_{918\text{-}1067\_L1047A\_L1050A}$ | 918-1067 aa (NdeI-XhoI) | Ni-NTA, TEV, Ni-NTA, Superdex 75 | ITC |

| | | | | | |
|---|---|---|---|---|---|
| | | | Quick change to mutate L1047 and L1050 to A | | |
| bMERB$_{W970A}$ | pET19mod-hs-Mical1_bMERB$_{W970A}$ | 6His-TEV-hs-Mical1$_{918\text{-}1067_W970A}$ | 918-1067 aa (NdeI-XhoI) Quick change to mutate W970 to A | Ni-NTA, TEV, Ni-NTA, Superdex 75 | ITC, aSEC |
| bMERB$_{V978A}$ | pET19mod-hs-Mical1_bMERB$_{V978A}$ | 6His-TEV-hs-Mical1$_{918\text{-}1067_V978A}$ | 918-1067 aa (NdeI-XhoI) Quick change to mutate V978 to A | Ni-NTA, TEV, Ni-NTA, Superdex 75 | ITC, aSEC, Crystallization |
| bMERB$_{V985A}$ | pET19mod-hs-Mical1_bMERB$_{V985A}$ | 6His-TEV-hs-Mical1$_{918\text{-}1067_V985A}$ | 918-1067 aa (NdeI-XhoI) Quick change to mutate V985 to A | Ni-NTA, TEV, Ni-NTA, Superdex 75 | ITC, aSEC, Crystallization |
| bMERB$_{V978A_V985A}$ | pET19mod-hs-Mical1_bMERB$_{V978A__V985A}$ | 6His-TEV-hs-Mical1$_{918\text{-}1067_V978A_V985A}$ | 918-1067 aa (NdeI-XhoI) Quick change to mutate V978 and V985 to A | Ni-NTA, TEV, Ni-NTA, Superdex 75 | ITC, aSEC, Crystallization |
| bMERB$_{L1011S}$ | pET19mod-hs-Mical1_bMERB$_{L1011S}$ | 6His-TEV-hs-Mical1$_{918\text{-}1067_L1011S}$ | 918-1067 aa (NdeI-XhoI) Quick change to mutate L1011 to S | Ni-NTA, TEV, Ni-NTA, Superdex 75 | aSEC |
| bMERB$_{V1038A}$ | pET19mod-hs-Mical1_bMERB$_{V1038A}$ | 6His-TEV-hs-Mical1$_{918\text{-}1067_V1038A}$ | 918-1067 aa (NdeI-XhoI) Quick change to mutate V1038 to A | Ni-NTA, TEV, Ni-NTA, Superdex 75 | aSEC |
| bMERB$_{V1041A}$ | pET19mod-hs-Mical1_bMERB$_{V1041A}$ | 6His-TEV-hs-Mical1$_{918\text{-}1067_V1041A}$ | 918-1067 aa (NdeI-XhoI) Quick change to mutate V1041 to A | Ni-NTA, TEV, Ni-NTA, Superdex 75 | aSEC |
| bMERB$_{I1048A}$ | pET19mod-hs-Mical1_bMERB$_{I1048A}$ | 6His-TEV-hs-Mical1$_{918\text{-}1067_I1048A}$ | 918-1067 aa (NdeI-XhoI) Quick change to mutate I1048 to A | Ni-NTA, TEV, Ni-NTA, Superdex 75 | aSEC |
| bMERB$_{V1041A_I1048A}$ | pET19mod-hs-Mical1_bMERB$_{V1041A_I1048A}$ | 6His-TEV-hs-Mical1$_{918\text{-}1067_V1ß41A_I1048A}$ | 918-1067 aa (NdeI-XhoI) Quick change to mutate V1041 and I1048 to A | Ni-NTA, TEV, Ni-NTA, Superdex 75 | ITC, aSEC |
| bMERB$_{L956S}$ | pET19mod-hs-Mical1_bMERB$_{L956S}$ | 6His-TEV-hs-Mical1$_{918\text{-}1067_L956S}$ | 918-1067 aa (NdeI-XhoI) Quick change to mutate L956 to S | Ni-NTA, TEV, Ni-NTA, Superdex 75 | ITC, aSEC |
| bMERB$_{S960D}$ | pET19mod-hs-Mical1_bMERB$_{S960D}$ | 6His-TEV-hs-Mical1$_{918\text{-}1067_S960D}$ | 918-1067 aa (NdeI-XhoI) Quick change to mutate S960 to D | Ni-NTA, TEV, Ni-NTA, Superdex 75 | ITC, aSEC |
| Rab8a$_{1\text{-}176}$ | pET19mod-hs-Rab8a$_{1\text{-}176opti}$ | Rai *et al.*, 2016[6] 6His-TEV-hs-Rab8a$_{1\text{-}176opti}$ | aa 1-176 codon-optimized (NdeI-XhoI) | Ni-NTA, TEV, Ni-NTA, Superdex75 | aSEC, ITC, Crystallization |
| Rab10$_{1\text{-}175}$ | pET19mod-hs-Rab10$_{1\text{-}175opti}$ | Rai *et al.*, 2016[6] 6His-TEV-hs-Rab10$_{1\text{-}175opti}$ | aa 1-175 codon-optimized (NdeI-XhoI) | Ni-NTA, TEV, Ni-NTA, Superdex75 | aSEC, ITC, Crystallization |
| Rab13$_{1\text{-}176}$ | pET19mod-hs-Rab13$_{1\text{-}176opti}$ | Rai *et al.*, 2016[6] 6His-TEV-hs-Rab13$_{1\text{-}176opti}$ | aa 1-176 codon-optimized (NdeI-XhoI) | Ni-NTA, TEV, Ni-NTA, Superdex75 | aSEC |

| | | | | | |
|---|---|---|---|---|---|
| $Rab15_{1-176}$ | pET19mod-hs-$Rab15_{1-176opti}$ | Rai *et al.*, 2016[6] 6His-TEV-hs-$Rab15_{1-176opti}$ | aa 1-176 codon-optimized (NdeI-XhoI) | Ni-NTA, TEV, Ni-NTA, Superdex75 | aSEC, ITC, |

aa: amino acid, hs: homo sapiens, fl: full-length

**Supplementary Table 3: Primers used in this study.**

| Primer name | Primer sequence (5´ to 3´) |
|---|---|
| Mical1_bMERB_R1044A FP | GTGGATTTGGTCAACCAGGCAGATGCCCTCATCCGCTTC |
| Mical1_bMERB_R1044A RP | GAAGCGGATGAGGGCATCTGCCTGGTTGACCAAATCCAC |
| Mical1_bMERB_L1011S FP | TGGCAGCTGGACCAGGAGAGTCGAGGCTACATGAACCGG |
| Mical1_bMERB_L1011S RP | CCGGTTCATGTAGCCTCGACTCTCCTGGTCCAGCTGCCA |
| Mical1_bMERB_V1038A FP | CAGGTCCTGAGGAAGCTGGCGGATTTGGTCAACCAGAGA |
| Mical1_bMERB_V1038A RP | TCTCTGGTTGACCAAATCCGCCAGCTTCCTCAGGACCTG |
| Mical1_bMERB_W970A FP | GAACAGCAAAAGAAACTAGCGGTAGGACAGCTGCTACAG |
| Mical1_bMERB_W970A RP | CTGTAGCAGCTGTCCTACCGCTAGTTTCTTTTGCTGTTC |
| Mical1_bMERB_V978A FP | GGACAGCTGCTACAGCTCGCTGACAAGAAAAACAGCCTG |
| Mical1_bMERB_V978A RP | CAGGCTGTTTTTCTTGTCAGCGAGCTGTAGCAGCTGTCC |
| Mical1_bMERB_V985A FP | GACAAGAAAAACAGCCTGGCGGCTGAGGAGGCCGAGCTC |
| Mical1_bMERB_V985A RP | GAGCTCGGCCTCCTCAGCCGCCAGGCTGTTTTTCTTGTC |
| Mical1_bMERB_V1041A FP | AGGAAGCTGGTGGATTTGGCCAACCAGAGAGATGCCCTC |
| Mical1_bMERB_V1041A RP | GAGGGCATCTCTCTGGTTGGCCAAATCCACCAGCTTCCT |
| Mical1_bMERB_I1048A FP | AACCAGAGAGATGCCCTCGCCCGCTTCCAGGAGGAGCGC |
| Mical1_bMERB_I1048A RP | GCGCTCCTCCTGGAAGCGGGCGAGGGCATCTCTCTGGTT |
| Mical1_bMERB_L956S FP | GTGAAGCTGGAGCTGGCCAGTAGGCGCCAGAGCAGTTCC |
| Mical1_bMERB_L956S RP | GGAACTGCTCTGGCGCCTACTGGCCAGCTCCAGCTTCAC |
| Mical1_bMERB_S960D FP | CTGGCCTTGAGGCGCCAGGACAGTTCCCCAGAACAGCAA |
| Mical1_bMERB_S960D RP | TTGCTGTTCTGGGGAACTGTCCTGGCGCCTCAAGGCCAG |
| Mical1_bMERB_L1047A_F1050A FP | GTCAACCAGAGAGATGCCGCCATCCGCGCCCAGGAGGAGCGCAGG |
| Mical1_bMERB_L1047A_F1050A RP | CCTGCGCTCCTCCTGGGCGCGGATGGCGGCATCTCTCTGGTTGAC |
| Mical1_bMERB_L974A_L975A FP | AAACTATGGGTAGGACAGGCGGCACAGCTCGTTGACAAGAAAAAC |
| Mical1_bMERB_L974A_L975A RP | GTTTTTCTTGTCAACGAGCTGTGCCGCCTGTCCTACCCATAGTTT |
| Mical1_bMERB_L977A_V978A FP | GTAGGACAGCTGCTACAGGCCGCTGACAAGAAAAACAGCCTGGTG |
| Mical1_bMERB_L977A_V978A RP | CACCAGGCTGTTTTTCTTGTCAGCGGCCTGTAGCAGCTGTCCTAC |
| Mical1_bMERB_M992A_I993A FP | GCTGAGGAGGCCGAGCTCGCGGCCACGGTGCAGGAATTGAATCTG |
| Mical1_bMERB_M992A_I993A RP | CAGATTCAATTCCTGCACCGTGGCCGCGAGCTCGGCCTCCTCAGC |
| Mical1_bMERB_E939A FP | CGGCGACTAAATGAGATTGCGGCTGCCTTGAGGGAGCTA |
| Mical1_bMERB_E939A RP | TAGCTCCCTCAAGGCAGCCGCAATCTCATTTAGTCGCCG |
| Mical1_bMERB_F925A FP | GAGGAGGAGATGAAGAGGGCCTGCAAGGCCCAGACCATC |
| Mical1_bMERB_F925A RP | GATGGTCTGGGCCTTGCAGGCCCTCTTCATCTCCTCCTC |
| Mical1_bMERB_K918A_E921A FP | TATTTTCAGGGCCATATGGCGGAGGAGGCGATGAAGAGGTTCTGC |
| Mical1_bMERB_K918A_E921A RP | GCAGAACCTCTTCATCGCCTCCTCCGCCATATGGCCCTGAAAATA |
